# NetCoMi: Network Construction and Comparison for Microbiome Data in R

**DOI:** 10.1101/2020.07.15.195248

**Authors:** Stefanie Peschel, Christian L. Müller, Erika von Mutius, Anne-Laure Boulesteix, Martin Depner

**Affiliations:** Institute for Asthma and Allergy Prevention, Helmholtz Zentrum München, German Research Center for Environmental Health, Munich, Germany; Department of Statistics, LMU München, Munich, Germany; Institute of Computational Biology, Helmholtz Zentrum München, German Research Center for Environmental Health, Munich, Germany; Center for Computational Mathematics, Flatiron Institute, New York, USA; Dr von Hauner Children’s Hospital, LMU München, Munich, Germany; Comprehensive Pneumology Center Munich (CPC-M), Member of the German Center for Lung Research, Munich, Germany; Institute for Medical Information Processing, Biometry and Epidemiology, LMU München, Munich, Germany

**Keywords:** compositional data, microbial association estimation, network analysis, sample similarity network, differential association, network comparison

## Abstract

Estimating microbial association networks from high-throughput sequencing data is a common exploratory data analysis approach aiming at understanding the complex interplay of microbial communities in their natural habitat. Statistical network estimation workflows comprise several analysis steps, including methods for zero handling, data normalization, and computing microbial associations. Since microbial interactions are likely to change between conditions, e.g. between healthy individuals and patients, identifying network differences between groups is often an integral secondary analysis step. Thus far, however, no unifying computational tool is available that facilitates the whole analysis workflow of constructing, analyzing, and comparing microbial association networks from high-throughput sequencing data.

Here, we introduce `NetCoMi` (**Net**work **Co**nstruction and comparison for **Mi**crobiome data), an `R` package that integrates existing methods for each analysis step in a single reproducible computational workflow. The package offers functionality for constructing and analyzing single microbial association networks as well as quantifying network differences. This enables insights into whether single taxa, groups of taxa, or the overall network structure change between groups. NetCoMi also contains functionality for constructing *differential networks*, thus allowing to assess whether single pairs of taxa are differentially associated between two groups. Furthermore, NetCoMi facilitates the construction and analysis of dissimilarity networks of microbiome samples, enabling a high-level graphical summary of the heterogeneity of an entire microbiome sample collection. We illustrate NetCoMi’s wide applicability using data sets from the GABRIELA study to compare microbial associations in settled dust from children’s rooms between samples from two study centers (Ulm and Munich).

**Availability:** A script with `R` code used for producing the examples shown in this manuscript are provided as Supplementary data. The NetCoMi package, together with a tutorial, is available at https://github.com/stefpeschel/NetCoMi.

## 1 Introduction

The rapid development of high-throughput amplicon sequencing techniques [1] offers new possibilities for investigating the microbiome across different habitats and provides the opportunity to discover relationships between the composition of microbial communities and their environment. Amplicon sequencing data are typically summarized in count tables where each entry expresses how often a read (a certain DNA or RNA sequence) associated with a specific taxon is observed in the sequencing process. Due to a lack of internal standards in common amplicon sequencing protocols, a particular feature of the data is that they carry relative or compositional information with each component expressing the relative frequency of a taxon in the sample [2]. Standard statistical analysis methods ignoring the compositional data structure may lead to spurious results (referred to as *compositional effects* in the following) [2]. In the following, we refer to approaches that take into account the compositional structure as *compositionally aware*.

Furthermore, the observed reads represent only a sample of the true microbial composition present in the biological material [3]. Accordingly, it is likely that not all taxa occurring in a sample are measured due to technical limitations in library preparation and the sequencing process [3]. Thus, the observed number of reads is only a noisy measurement reflecting the probability of the corresponding organisms to be present [2]. Moreover, amplicon sequencing data collections comprise a high amount of zero counts, contain samples with varying sequencing depths (sum of counts per sample), and are usually high dimensional, i.e., the number of taxa *p* is much higher than the sample size *n*.

A common exploratory analysis approach for microbiome survey data is the estimation of microbe-microbe association networks [4], allowing for high-level insights into the global structure of microbial communities. Existing approaches for measuring and estimating microbial associations include compositionally aware correlation estimators, such as SparCC [5], partial correlation estimators [6, 7], and proportionality [8]. A wide range of existing general-purpose tools is available for visualizing and analyzing networks. Popular `R` packages are `igraph` [9], `statnet` [10], and `network` [11]. Software packages such as Gephi [12] and Cytoscape [13] provide functionalities for large-scale network visualization, analysis, and network generation. However, a naive application of these tools on compositional data bears the risk of resulting in a network that contains spurious associations.

While the analysis of a single microbial association network can provide insights into the general organizational structure of a microbial community, researchers are often more interested in how microbial associations *change* across different conditions. For instance, it is often desired to find relationships between microbial compositions, their inherent connectivity, and an underlying phenotype, e.g. the health status of patients. This task thus requires the quantitative comparison of networks across conditions.

Current approaches for comparing networks between two conditions can be divided into two types: (i) *differential association analysis* focusing on differences in the strength of single associations, and (ii) *differential network analysis*, analyzing differences between network metrics and network structure between two conditions [14]. Differential associations can further be used as the basis for constructing differential networks, where only differentially associated nodes are connected (see [14] for a comparative study of differential network analysis methods). Existing tools for network comparison either require pre-computed networks (adjacency matrix or edge list) as input (e.g. CompNet [15]) or are tailored for protein interaction (e.g. NetAlign [16] and Netdis [17]) or gene functions [18], not for microbiome data.

In this paper, we introduce `NetCoMi` (**Net**work **Co**nstruction and comparison for **Mi**crobiome data), a comprehensive `R` package, that integrates previously disjoint microbial network inference and analysis tasks into a single coherent computational workflow. `NetCoMi` allows the user to construct, analyze, and compare microbial association networks in a fast and reproducible manner. The complete `NetCoMi` workflow is shown in Fig. 1. `NetCoMi` provides a wide range of existing methods for data normalization, zero handling, edge filtering, and a selection of association measures, which can be combined in a modular fashion to generate the microbial networks. For network analysis, several local and global network properties are provided, which can be visualized in network plots to enable a descriptive comparison. Quantitative comparison between two different networks is available via the integration of appropriate statistical tests. Furthermore, our package enables (i) the generation of differential microbial networks and (ii) the construction of *sample similarity networks* (using, e.g. the Bray-Curtis measure), which can serve as a high-level visual summary of the heterogeneity of the microbiome sample collection.

**Fig. 1.**
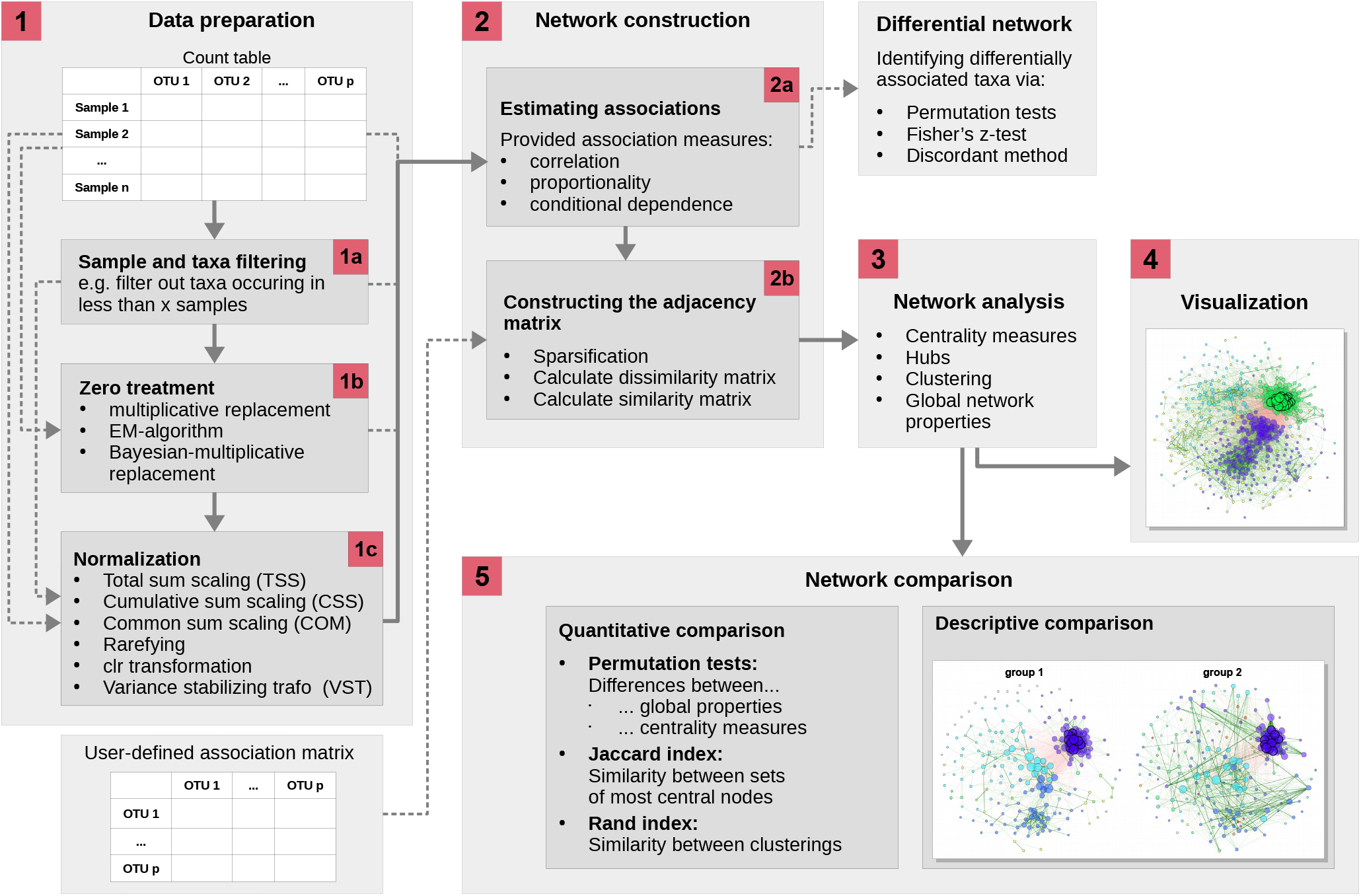
The proposed workflow for constructing, analyzing, and comparing microbial association networks, implemented in the `R` package `NetCoMi`. The main framework (displayed as continuous lines) requires a *n × p* read count matrix as input. The data preparation step includes sample and taxa filtering, zero replacement, and normalization (step 1). Associations are calculated and stored in an adjacency matrix (step 2). Alternatively, an association matrix is accepted as input, from which the adjacency matrix is determined. A more detailed chart describing step 2 is given in Fig. 2. In step 3, network metrics are calculated, which can be visualized in the network plot (step 4). If two networks are constructed (by passing either a binary group vector or two user-defined association matrices to the function), their properties can be compared (step 5). Besides the main workflow, a differential network can be constructed from the association matrix.

**Fig. 2.**
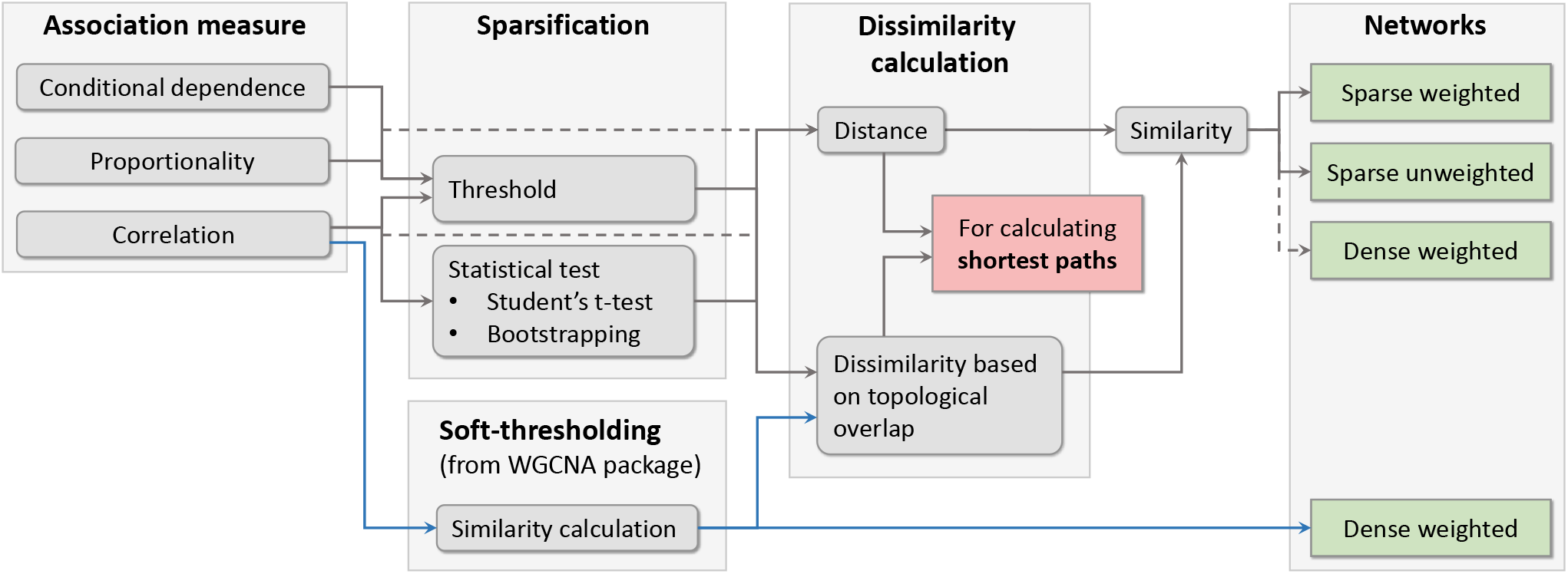
Approaches for network construction that are available in `NetCoMi`, depending on the association measure. For correlations, in addition to a threshold and statistical testing, the soft-thresholding approach from `WGCNA` package [35] is implemented (marked by blue arrows). Dissimilarity based on topological overlap (also adopted from `WGCNA` package) is available as a further dissimilarity transformation approach in addition to metric distances and thus used for all network properties based on shortest paths. Whether a network measure is based on similarity or dissimilarity is stated in Table 4. Network construction without a sparsification step leads to dense networks where all nodes are connected.

## 2 Network construction and characterization

### 2.1. Data filtering, normalization, and zero handling

The process of constructing microbial association networks starts with a matrix containing absolute read counts originating from a sequencing process. The total read counts 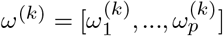 of a sample *k* with *p* taxa are a composition summing up to a constant 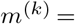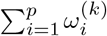, the sequencing depth. The sequencing depth differs from sample to sample and is predefined by technical factors leading to sparse data with many zeros. Thus, preprocessing steps (step 1 in Fig. 1)are recommended, or even mandatory depending on the association measure (see Supplementary Table S1).

To simplify the graphical interpretation of an association network and the computational processes, it is reasonable to filter out a certain set of taxa as first data preparation step (step 1a in Fig. 1). See Table S2 for the options available in `NetCoMi`.

The excess number of zeros in the data is a major challenge for analyzing microbiome data because parametric as well as non-parametric models may become invalid for data with a large amount of zeros [22]. Moreover, many compositionally aware measures are based on so-called log-ratios. Log-ratios have been proposed by Aitchison [28] as the basis for statistical analyses of compositional data as they are independent of the total sum of counts *m*. More precisely, for two variables *i* and *j* the log-ratio of relative abundances 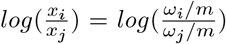 is equal to the log-ratio of the absolute abundances 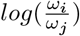 [6]. However, log-ratios cannot be computed if the count matrix contains any zeros, making zero handling necessary (step 1b in Fig. 1). Several zero replacement strategies have been proposed [24, 23, 25, 29, 27]. Table 1 gives an overview of the different types of zeros that have been suggested as well as existing approaches for their treatment.

**Table 1.** Different zero types defined for sequencing data with appropriate approaches for their treatment (suggested by Martín-Fernández et al. [19]), which are available in `NetCoMi`. The three approaches are implemented in R via the zCompositions package [20]. Shown are for each treatment method a few key facts as well as the argument for applying it in `NetCoMi`.

| Type | essential/structural zeros | rounded zeros | count zeros |
| --- | --- | --- | --- |
| Description | “...component which is truly zero, not something recorded as zero simply because the experimental design or the measuring instrument has not been sufficiently sensitive to detect a trace of the part.” [21] | Assumed to be present in the sample but their proportion is too low to be detected [19]. | Sequencing data are seen as categorical. For a zero count it is assumed that, in this category, no event occurred in this experimental run but could occur in another run (e.g. with a different sequencing depth) [19]. |
| Treatment methods | No general approach available so far [22]<br>The “structural zero issue is by far the most complicated problem” [22]<br>⇒ no approach included in NetCoMi | Multiplicative imputation [23]:<br>Non-parametric<br>Small value $\delta$ imputed for all zeros<br>Non-zero values adjusted regarding unit-sum constraint to preserve their covariance structure<br>NetCoMi arg.: `zeroMethod = "multRepl"`<br>Modified EM *alr*-algorithm [24, 25, 26]:<br>Parametric<br>Covariance structure is taken into account<br>Zeros imputed via EM-algorithm for *alr* transformed data<br>NetCoMi arg.: `zeroMethod = "alrEM"` | Bayesian-multiplicative treatment [27]:<br>Parametric<br>Observed counts $\mathbf{c} = (c_1, \dots, c_p)$ assumed to be categorical<br>Random vector $\mathbf{C}$ multinomial distributed with parameters $(n, \pi_1, \dots, \pi_p)$<br>$\pi$ corresponds to observed microbial proportions → estimated via posterior distribution<br>NetCoMi arg.: `zeroMethod = "bayesMult"` |

Normalization techniques are required to make read counts comparable across different samples [30, 31] (step 1c in Fig. 1). The normalization approaches included in `NetCoMi` are summarized in Table 2. A description of these methods is available in [30] and [31]. Note that forcing the read counts of each sample to a unique sum (as done with Total sum scaling) does not change the compositional structure. Rather, proportions are always compositional, even if the original data are not [2]. Aitchison [28] suggested using the centered log-ratio (clr) transformation to move compositional data from the simplex to real space. Badri et al. [30] have shown that variance-stabilizing transformations (VSTs) [32] as well as the clr-transformation produce very similar Pearson correlation estimates, which are more consistent across different sample sizes than TSS, CSS, and COM methods.

**Table 2.**
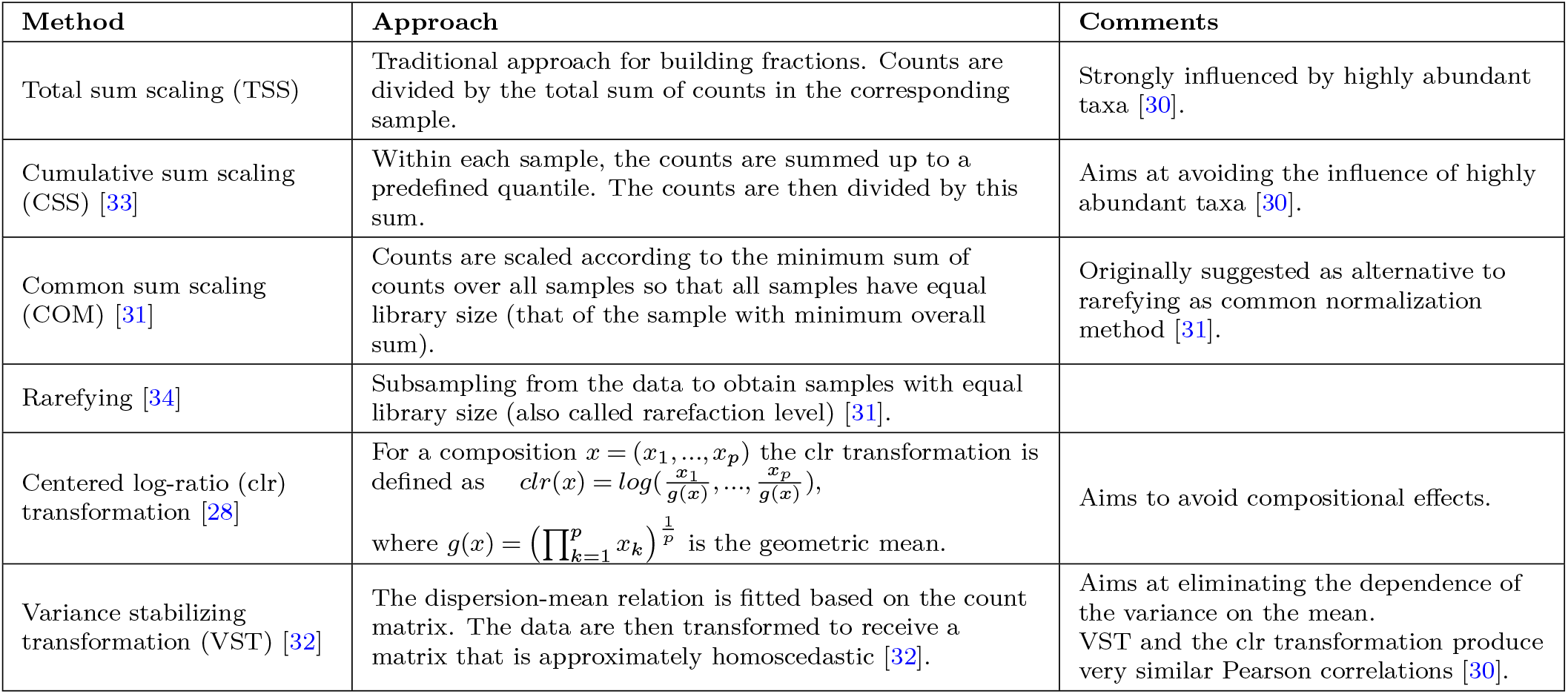
Data normalization techniques implemented in `NetCoMi`. More detailed descriptions of these methods can be found in Badri et al. [30] and McMurdie and Holmes [31]. The corresponding `NetCoMi` argument is normMethod with the available options: “none”, “TSS”, “CSS”, “COM”, “rarefy”, “clr”, and “VST”.

| Method | Approach | Comments |
| --- | --- | --- |
| Total sum scaling (TSS) | Traditional approach for building fractions. Counts are divided by the total sum of counts in the corresponding sample. | Strongly influenced by highly abundant taxa [30]. |
| Cumulative sum scaling (CSS) [33] | Within each sample, the counts are summed up to a predefined quantile. The counts are then divided by this sum. | Aims at avoiding the influence of highly abundant taxa [30]. |
| Common sum scaling (COM) [31] | Counts are scaled according to the minimum sum of counts over all samples so that all samples have equal library size (that of the sample with minimum overall sum). | Originally suggested as alternative to rarefying as common normalization method [31]. |
| Rarefying [34] | Subsampling from the data to obtain samples with equal library size (also called rarefaction level) [31]. |  |
| Centered log-ratio (clr) transformation [28] | For a composition $x = (x_1, \dots, x_p)$ the clr transformation is defined as $clr(x) = \log(\frac{x_1}{g(x)}, \dots, \frac{x_p}{g(x)})$, where $g(x) = (\prod_{k=1}^p x_k)^{\frac{1}{p}}$ is the geometric mean. | Aims to avoid compositional effects. |
| Variance stabilizing transformation (VST) [32] | The dispersion-mean relation is fitted based on the count matrix. The data are then transformed to receive a matrix that is approximately homoscedastic [32]. | Aims at eliminating the dependence of the variance on the mean. VST and the clr transformation produce very similar Pearson correlations [30]. |

### 2.2. Measuring associations between taxa

Association estimation is the next step in our workflow (step 2a in Fig. 1) to obtain statistical relations between the taxa. Common association measures include correlation, proportionality, and conditional dependence. For all three types of association, compositionally aware approaches have been proposed, which are summarized in Table 3. Further information on these measures is available in Supplement 1. To ensure wide applicability of `NetCoMi`, the package comprises also traditional association measures, which are not suitable for application on read count data in their original form.

**Table 3.** Overview of compositionally aware association measures that are available for network construction in `NetCoMi`. The corresponding argument is named measure. The measures are grouped by the three types of association: correlation, proportionality and conditional dependence. For each measure, the assumptions stated in the corresponding publication are listed, together with a short summary of the approach.

| Measure | Data transformation<br>R implementation | Assumptions | Approach |
| --- | --- | --- | --- |
| Correlation |  |  |  |
| SparCC [5] | Log-ratios<br>R code based on package `r-sparcc` [44] | Large number of taxa<br>Sparse underlying network (taxa sparsely correlated) | Idea: Estimating Pearson correlations from log-ratio variances via an approximation based on the assumption that taxa are uncorrelated on average<br>Zero handling: Bayesian approach (observations replaced by random samples)<br>Nested iterations to reinforce sparsity assumption and account for uncertainty due to random sampling |
| CCLasso [36] | Log-ratios<br>R code on GitHub [45] | Large number of taxa<br>Sparse underlying network | Idea: Inferring correlations from observed counts via latent variable model<br>Approach: minimizing a loss function (considers errors resulting from estimating the covariance matrix from observed data instead of true abundances) plus a penalty to incorporate the sparsity assumption |
| CCREPE [37] | Relative abundances<br>R package `ccrepe` [46] | — | Idea: Testing for statistical significance of the estimated correlations, whereby spurious correlations caused by compositional effects are considered<br>Approach: permuting and re-normalizing the data to assess correlations due to compositionality alone |
| Proportionality |  |  |  |
| $\rho$ [38] | clr transformation<br>R package `propr` [8] | — | $\rho(\log x, \log y) = 1 - \frac{\text{var}(\log(x/y))}{\text{var}(\log x) + \text{var}(\log y)}$ |
| Conditional dependence |  |  |  |
| SPIEC-EASI [47] | Log-ratios<br>R package `SpiecEasi` [47] | Large number of taxa<br>Sparse underlying network | Idea: covariance matrix for clr-transformed data is approximately equal to the “true” covariance matrix $\Sigma$ for $p \gg 0$<br>Zero entries in the inverse covariance matrix $\Sigma^{-1}$ correspond to conditional independent taxa<br>Approach: a graphical model is used to infer the conditional dependence structure between each two taxa<br>Two methods for graphical model inference: sparse inverse covariance selection and neighborhood selection |
| gCoda [40] | Relative abundances<br>R code on GitHub [48] | True absolute abundances $y$ follow a multivariate normal distribution<br>Sparse underlying network | $\log(y) \sim N(\mu, \Sigma)$ with $\Omega = \Sigma^{-1}$ representing conditional dependence relationships between taxa<br>$\Omega$ is determined by minimizing the negative log-Likelihood for $(\mu, \Omega)$ plus $l_1$ penalty (to satisfy sparsity assumption)<br>Optimization problem solved via Majorization- Minimization algorithm |
| SPRING [39] | *mclr* transformation<sup>1</sup><br>R package `SPRING` [49] | Sparse underlying network | $\Sigma$ is estimated using a semi-parametric rank based approach relying on a truncated Gaussian copula model [50], which can deal with zeros in the data<br>Conditional independence relationships are inferred from $\Sigma$ using neighborhood selection [42]<br>Graphical model selection via stability-based approach |

<sup>1</sup> *clr* transformation where only non-zero elements are included in the geometric mean [39].

#### 2.2.1. Correlations

Compositionality implies that if the absolute abundance of a single taxon in the sample increases, the perceived relative abundance of all other taxa decreases. Applying traditional correlation measures such as Pearson’s correlation coefficient to compositions can lead to spurious negative correlations, which do not reflect underlying biological relationships. It has been shown that the resulting bias is stronger, the lower the diversity of the data [5].

A possible approach to reduce compositional effects is to normalize or transform the data in a compositionally aware manner and apply standard correlation coefficients to the transformed data. A common method is the clr transformation (see Table 2), which is a variation of the aforementioned log-ratio approach. Traditional correlation measures provided in `NetCoMi` are the Pearson correlation, Spearman’s rank correlation, and the biweight midcorrelation (bicor). Bicor as part of the `R` package `WGCNA` [35] is more robust to outliers than Pearson’s correlation because it is based on the median instead of the mean of observations.

Popular compositionally aware correlation estimators include SparCC (Sparse Correlations for Compositional data) [5], CCLasso (Correlation inference for correlations for compositional data and has become Compositional data through Lasso) [36], and CCREPE (Compositionality Corrected by REnormalization and PErmutation), also called ReBoot method [37]. SparCC is one of the first approaches developed for inferring correlations for compositional data and has become a widely-used method [36]. However, SparCC has some limitations, namely that the estimated correlation matrix is not necessarily positive definite and may have values outside [−1,1] [36]. Furthermore, the basic algorithm is repeated iteratively to reinforce the sparsity assumption and to account for uncertainties due to random sampling, leading to high computational complexity. CCLasso is also based on log-ratios but aims to avoid the disadvantages of SparCC by using a latent variable model to infer a positive definite correlation matrix directly [36]. The CCREPE approach operates directly on relative read counts, which are permuted and re-normalized in order to detect correlations induced by compositionality alone.

#### 2.2.2. Proportionality

Lovell et al. [38] argue that correlations cannot be inferred from the relative abundances in a compositionally aware manner without any assumptions and propose proportionality as an alternative association measure for compositional data. If two relative abundances are proportional, then their corresponding absolute abundances are proportional as well: 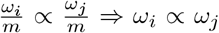. Thus, proportionality is identical for the observed (relative) read counts and the true unobserved counts. Lovell et al. [38] suggest proportionality measures based on the log-ratio variance 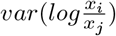, which is zero when *ω_i_* and *ω_j_* are perfectly proportional. This variance, however, lacks a scale that would make the strength of association comparable. For this reason, the proposed proportionality measures *ϕ* and *ρ* [38, 8] are modifications of the log-ratio variance based on clr transformed data that come with a scale. Due to its analogy to correlations, we included *ρ* in the `NetCoMi` package (see Table 3 for the formula). *ρ* is a symmetric measure with values in [−1,1], where 1 corresponds to perfect proportionality.

#### 2.2.3. Conditional dependence

Conditional dependence expresses the relation between two variables conditioned on all other variables in the data set [6]. Hence, this is a measure of direct relations between each two taxa, while (marginal) correlations cannot differentiate between direct and indirect dependencies. Three estimators of conditional dependence are included in `NetCoMi`: SPRING (Semi-Parametric Rank-based approach for INference in Graphical model) [39], SPIEC-EASI (Sparse Inverse Covariance Estimation for Ecological Association Inference) [6], and gCoda [40].

All three approaches use graphical models to infer the conditional independence structure from the data. For multivariate Gaussian data, the graph structure can be inferred from the non-zero elements of the inverse covariance matrix Ω = Σ^−1^ [41], where each entry is related to scaled negative partial correlation. Loh and Wainwright [41] relaxed the Gaussianity assumption and established relationships between the inverse covariance matrix and the edges of a graph for discrete data. All three approaches for estimating a conditional dependence graph presume that the assumptions of the data generation process are fulfilled so that the graph structure can be reliably inferred from the count matrix. We also consider the assumptions on the data generation process as satisfied and use *conditional dependence* and *partial correlation* equivalently in the following. gCoda uses a Majorization-Minimization algorithm to infer Ω from the data based on maximizing a penalized likelihood. In SPIEC-EASI, two approaches are provided to obtain Ω from the observed count data: neighborhood selection [42] and sparse inverse covariance selection (also known as “graphical lasso”) [43]. SPRING uses a semi-parametric rank-based correlation estimator which can account for excess zeros in the data and applies neighborhood selection to infer conditional dependencies. All three measures assume a sparsely connected underlying network.

### 2.3. Constructing the adjacency matrix

Using one of the aforementioned association measures, an association matrix with entries *r*_*ij*_ expressing the relation between pairs of taxa *i* and *j* is computed. The next step is sparsification and transformation into distances and similarities (step 2b in Fig. 1) resulting in an adjacency matrix *A*^*p×p*^ with entries *a*_*ij*_ as numerical representation of the microbial network, where nodes (or vertices) represent the taxa. The different options available in `NetCoMi` are illustrated in Fig. 2 (see Supplementary Figure S1 for a more detailed version of this chart).

Since the estimated associations are generally different from zero, using them directly as adjacency matrix results in a *dense network*, where all nodes are connected to each other and consequently only weighted network measures are meaningful. Instead, the association matrix is usually sparsified to select edges of interest. One possible sparsification strategy consists of defining a cutoff value (or threshold) so that only taxa with an absolute association value above this threshold are connected [5, 51]. This filtering method is available for all types of association. The conditional independence measures SPRING, SPIEC-EASI, and gCoda already include a model selection approach, making the filter step unnecessary.

For correlations, statistical tests are available as alternative sparsification method, allowing only significant associations to be included in the network. Student’s t-test [52] and a bootstrap approach [5, 51] are implemented for identifying correlations significantly different from zero. The resulting p-values need to be adjusted to account for multiple testing. `NetCoMi` includes all adjustment methods available in the `R` function `p.adjust()` (`stats` [53]) and, in addition, two methods for multiple testing under dependence having higher power than the common Benjamini-Yekutieli method [54]: (i) `fdrtool()` for controlling the local false discovery rate [55, 56], and (ii) the adaptive Benjamini-Hochberg method [57] where the proportion of true null hypotheses is estimated using convex decreasing density estimation as proposed by Langaas et al. [58].

P-values arising from bootstrapping might be problematic because they can be exactly zero if none of the bootstrap-correlations is more extreme than the observed one. Since p-values of exactly zero cannot be corrected for multiple testing, `NetCoMi`’s bootstrap p-values are corrected by adding pseudo counts:

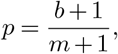

where *b* is the number of generated samples with a test statistic at least as extreme as the observed one, and *m* is the number of repetitions [59].

As illustrated in Figure S1, the sparsified associations 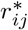 are then transformed to dissimilarities/distances *d*_*ij*_, which are needed for computing network measures based on shortest paths (see Section 2.4). Following van Dongen and Enright [60], we included different distance metrics depending on how negative associations should be handled (see Figure S1). Available are the options: (i) “unsigned”: 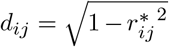, leading to a low distance between strongly associated taxa (positively as well as negatively), (ii) “signed”: 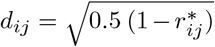, where the distance is highest for strongly negative associated taxa, and (iii) “signedPos”: “signed” distance with setting negative associations to zero. A dissimilarity measure based on the topological overlap matrix (TOM) [61] is also available in `NetCoMi`.

The adjacency matrix contains similarities of the form *s*_*ij*_ = 1 − *d*_*ij*_. These similarity values are used for network plot and network metrics based on connection strength (see Section 2.4). `NetCoMi` also offers two options for constructing an unweighted network via generating a binary adjacency matrix from the sparse association matrix where the user can decide whether or not negative associations should be included in the unweighted network (Figure S1).

Furthermore, we included adjacency matrix constructions via the *soft-thresholding* approach from the `WGCNA` package [35, 29], which is only available for correlations (see blue path in Fig. 2). Rather than using *hard thresholding* (e.g. via cutoff value or statistical tests), Zhang and Horvath [29] suggested raising the estimated correlations to the power of a predefined value greater than 1 to determine the adjacencies (see Supplement 2.2 for more details). Small correlation values are thus pushed toward zero becoming less important in the network. The resulting similarities are used as edge weights for network plotting and network metrics based on connection strength. Note that this approach leads to a fully connected network. Following Zhang and Horvath [29], only TOM dissimilarities are available in `NetCoMi` for computing shortest paths and clusters when soft-thresholding is used.

### 2.4. Network analysis

We analyze a constructed network by calculating network summary metrics (step 3 in Fig. 1), which are amenable to group comparisons. Alternatively, `NetCoMi` offers the possibility to analyze and visualize single networks.

Several network statistics require shortest path calculations. A *path* between two vertices *v*_0_ and *v*_*k*_ is a sequence of edges connecting these vertices such that *v*_0_ and *v*_*k*_ are each at one end of the sequence and no vertices are repeated [62]. The *length* of a path is the sum of edge weights, where the weight of an edge is a real non-negative number associated with this edge [62]. The *shortest path* between two nodes is the path with minimum length. In `NetCoMi`, edge weights are defined in two ways: (i) For properties based on shortest paths, dissimilarity values are used implying that the path length between two taxa is shorter the higher the association between the taxa is. (ii) For properties based on connection strength, the corresponding similarities are used as edge weights (see Section 2.3 and Figure S1 for details on distance and similarity calculation).

Network centrality measures offer insights into the role of individual taxa within the microbial community. We consider degree, betweenness, closeness, and eigenvector centrality (see Table 4). Using these measures, so-called *hubs* (or keystone taxa) can be determined. This analysis step is of high interest to researchers since hub nodes may correspond to taxa with a particularly important role in the microbial community. What is deemed “important”, depends on the scientific context. A selection of definitions for hub nodes is given in Table 4.

**Table 4.** Local and global network properties implemented in `NetCoMi`. The table shows for each measure a short explanation and whether it is based on connection strength (similarity) or dissimilarity/distance (e.g. two taxa with a high correlation have a high connection strength, but their distance is low).

| Measure | Definition | Based on |
| --- | --- | --- |
| Shortest Path | Sequence of edges connecting two nodes with minimum sum of edge weights [62]. | Dissimilarity |
| **Local measures (refer to single nodes)** |  |  |
| Degree centrality | Number of adjacent nodes [63]. Options: weighted/unweighted. | Similarity |
| Betweenness centrality | Fraction of times a node lies on the shortest path<sup>1</sup> between all other nodes [63].<br>⇒ A central node has the ability to connect sub-networks [51]. | Dissimilarity |
| Closeness centrality | Reciprocal of the sum of shortest paths between this node and all other nodes [63].<br>⇒ The node with highest closeness centrality has the minimum shortest path to all other nodes. | Dissimilarity |
| Eigenvector centrality | Calculated via eigenvalue decomposition: $Ac = \lambda c$, where $\lambda$ denotes the eigenvalues and $c$ the eigenvectors of the adjacency matrix $A$. Eigenvector centrality is then defined as the $i$-th entry of the eigenvector belonging to the largest eigenvalue [64, 65].<br>⇒ A node is central if it is connected to other nodes having themselves a central position in the network. | Similarity |
| Normalized centralities<sup>3</sup> | For each centrality measure, a normalized version leading to values in [0,1] is implemented in NetCoMi. We use the following definitions of normalized centrality for a vertex $v_i$:<br>Degree: $C_{\text{Deg}}^*(v_i) = \frac{1}{n-1} C_{\text{Deg}}(v_i)$ [63],<br>Betweenness centrality: $C_{\text{Betw}}^*(v_i) = \frac{2}{n^2-3n+2} C_{\text{Betw}}(v_i)$ [63],<br>Closeness centrality: $C_{\text{Close}}^*(v_i) = \frac{1}{n-1} C_{\text{Close}}(v_i)$ [63],<br>Eigenvector centrality: $C_{\text{Eigen}}^*(v_i) = \frac{C_{\text{Eigen}}(v_i)}{\max(C_{\text{Eigen}}(v))}$ [66],<br>where $n$ is the number of nodes in the network, respectively. | |
| Hubs | Particularly important nodes, which are most central regarding:<br>1) highest degree centrality [67],<br>2) highest degree, betweenness and closeness centrality at the same time [68],<br>3) or highest eigenvector centrality [9].<br>For each centrality measure, the *most central nodes* are those with a centrality value either above a certain quantile of the fitted log-normal distribution [68] or above a certain empirical quantile. | Depends on the centrality measure. |
| **Global network metrics (refer to the whole network)** |  |  |
| Average path length | Arithmetic mean of all shortest paths between vertices in a network [67]. | Dissimilarity |
| Global clustering coefficient | Proportion of triangles with respect to the total number of connected triples<sup>2</sup> [67].<br>⇒ Expresses how likely the nodes are to form clusters [67].<br>For weighted networks, the definition according to Barrat et al. [69] is used in NetCoMi. | Similarity |
| Modularity | Expresses how well the network is divided into communities (many edges within the identified clusters and only a few between them) [70]. | Depends on the clustering algorithm |
| Edge / vertex connectivity | Minimum number of edges, or vertices (nodes) that need to be removed to disconnect the network, respectively [71]. Not meaningful for a fully connected network. | Presence/absence of an edge |
| Density | Ratio of the actual number of edges in the network and the possible number of edges [71]. Not meaningful for a fully connected network. | Presence/absence of an edge |
| **Clustering** |  |  |
| Hierarchical clustering | Hierarchical clustering using the R function `hclust()` from `stats` package [53]. Different methods are provided (e.g. single, complete, and average linkage, Ward's method). The `cutree()` function (`stats`) is used for cutting the resulting tree. | Dissimilarity |
| Modularity clustering [72] | The modularity measure is maximized over all possible clusterings. | Similarity |
| Fast greedy modularity optimization [70] | Modularity clustering via a fast greedy algorithm, which is suitable for very large networks. | Similarity |
| Clustering based on edge betweenness [73] | Idea: Edges with high edge betweenness (number of shortest paths leading through an edge) tend to divide the network into clusters. Approach: Hierarchical clustering, where edges with high edge betweenness are removed gradually from the network. | Dissimilarity |

<sup>1</sup> The shortest path is given as the path between two nodes that minimizes the sum of distances between included nodes.
<sup>2</sup> A connected triple is a group of three nodes with two or three edges. A triangle is a triple where all three nodes are connected.
<sup>3</sup> Not to be confused with “centralization” [63], which is a global network measure expressing the degree to which the centrality of the most central node exceeds the centrality of all other nodes in the network .

Clustering methods are appropriate to identify functional groups within a microbial community. A *cluster* (or module) is a group of nodes that are highly connected to one another but have a small number of connections to nodes outside their module [51]. The user of `NetCoMi` can choose between one of the clustering methods provided by the `igraph` package [9] and hierarchical clustering (`R` package `hclust()` from `stats` package). Both similarities and dissimilarities can be used for clustering (see Table 4).

Global network properties are defined for the whole network and offer an insight into the overall network structure. Typical measures are the average path length, edge and vertex connectivity, modularity, and the clustering coefficient (see Table 4).

### 2.5. Sample similarity networks

Using dissimilarity between samples rather than association measures among taxa leads to networks where nodes represent subjects or samples rather than taxa. Analogous to microbiome ordination plots, these *sample networks* express how similar microbial compositions between subjects are and thus provide insights into the global heterogeneity of the microbiome sample collection.

The following dissimilarity measures are available in `NetCoMi`: Euclidean distance, Bray-Curtis dissimilarity [74], Kullback-Leibler divergence (KLD) [75], Jeffrey’s divergence [76], Jensen-Shannon divergence [77], compositional KLD [78, 79], and Aitchison distance [80]. Details are available in Table S3. Only Aitchison’s distance and the compositional KLD are suitable for application on amplicon sequencing data while the others may induce compositional effects when applied to raw count data without appropriate transformation (see Table 2).

The workflow for constructing, analyzing, and comparing sample similarity networks is described in Supplement 2.5. For these networks, the same network properties as for microbial networks are available in `NetCoMi` (Table 4). While the estimated dissimilarity values *d_kl_* between two samples *k* and *l* are used for network properties based on shortest paths, the corresponding similarities, calculated by

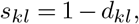

are used for properties based on connection strength and edge weights in the network plot. Accordingly, highly connected nodes are subjects with a microbial composition similar to many other subjects. Furthermore, as in usual cluster analysis, clusters represent subjects with similar bacterial composition but with the advantage, that the solution is visualized in the network plot.

## 3 Network comparison

`NetCoMi`’s network comparison module (step 5 in Fig. 1) focuses on investigating the following questions in a quantitative fashion: (i) Is the overall network structure different between two groups? (ii) Are hub taxa different between the two microbial communities? (iii) Do the microorganisms build different “functional” groups? (iv) Are single pairs of taxa differentially associated among the groups? To meet these objectives, `NetCoMi` offers several network property comparison modes as well as the estimated associations itself between two groups. These approaches are commonly used in other fields of application and have been adapted to the microbiome context. Each method includes statistical tests for significance.

To perform network comparison, the count matrix is split into two groups according to the user-defined group indicator vector. All steps of network construction and analysis described in Section 2 are performed for both subsets separately. The estimated associations, as well as network characteristics, are then compared using the methods described below.

### 3.1. Differential network analysis

We next detail `NetCoMi`’s differential network analysis capabilities. All approaches are applicable for microbial association networks and sample similarity networks, respectively.

#### 3.1.1. Permutation tests

`NetCoMi` uses permutation tests to assess for each centrality measure and taxon whether the calculated centrality value is significantly different between the two groups. The null hypothesis of these tests is defined as 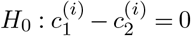, where 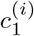 and 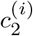 denote the centrality value of taxon *i* in group 1 and 2, respectively. A standard non-parametric permutation procedure [81] is used to generate a sampling distribution of the differences under the null hypothesis (see Supplement 3.1). The same approach is used to test for significant group differences in the global network characteristics listed in Table 4.

#### 3.1.2. Similarity of most central nodes

A set of *most central nodes* can be defined in two ways: In the first approach, under the assumption that the centrality values are log-normal distributed [68], the set of most central nodes contains nodes with a centrality value greater than a certain quantile of the fitted log-normal distribution. In the second approach, the empirical quantile can be used directly without any distributional assumption. Both approaches are included in `NetCoMi`, where the quantile can be freely chosen in each case.

The Jaccard index [82] (see Supplement 3.2 for a definition) can then be used for assessing h ow different the two sets of most central nodes (regarding a certain centrality measure) are between the groups. This index ranges from zero to one, where a value of one corresponds to two equal sets and zero means that the sets have no members in common. Following Real and Vargas [83], we included an approach to test whether the observed value of Jaccard’s index is significantly different from that expected at random (see Supplement 3.2). Note that this approach cannot make a statement on whether the two sets of most central nodes are significantly different.

#### 3.1.3. Similarity of clustering solutions

`NetCoMi` offers several network partitioning and clustering algorithms (see Table 4). One way to assess the agreement of two partitions is via the Rand index [84] (see Supplement 3.3 for a definition). Like Jaccard’s index, the R and index ranges from zero to one, where one indicates that the clusters are exactly equal in both groups. The original Rand index is dependent on the number of clusters making it difficult to interpret. Instead, `NetCoMi` uses the adjusted Rand index [85]. The adjusted values take values in the interval [−1, 1], where one corresponds to identical clusterings and zero to the expected value for two random clusterings. Consequently, positive index values imply that two clusterings are more similar and negative values less similar than expected at random. Following [86], `NetCoMi` uses a permutation procedure to test whether a calculated value is significantly different from zero. However, this test does not signify whether the clusterings are significantly different between the groups. Details about this approach and its implementation in R are given in Supplement 3.3.

### 3.2. Differential association analysis

`NetCoMi` also allows to test differences between the estimated associations themselves rather than network properties. This analysis is referred to as *differential association analysis*. Table 5 shows the three approaches available in `NetCoMi` (see Supplement 4 for details).

**Table 5.**
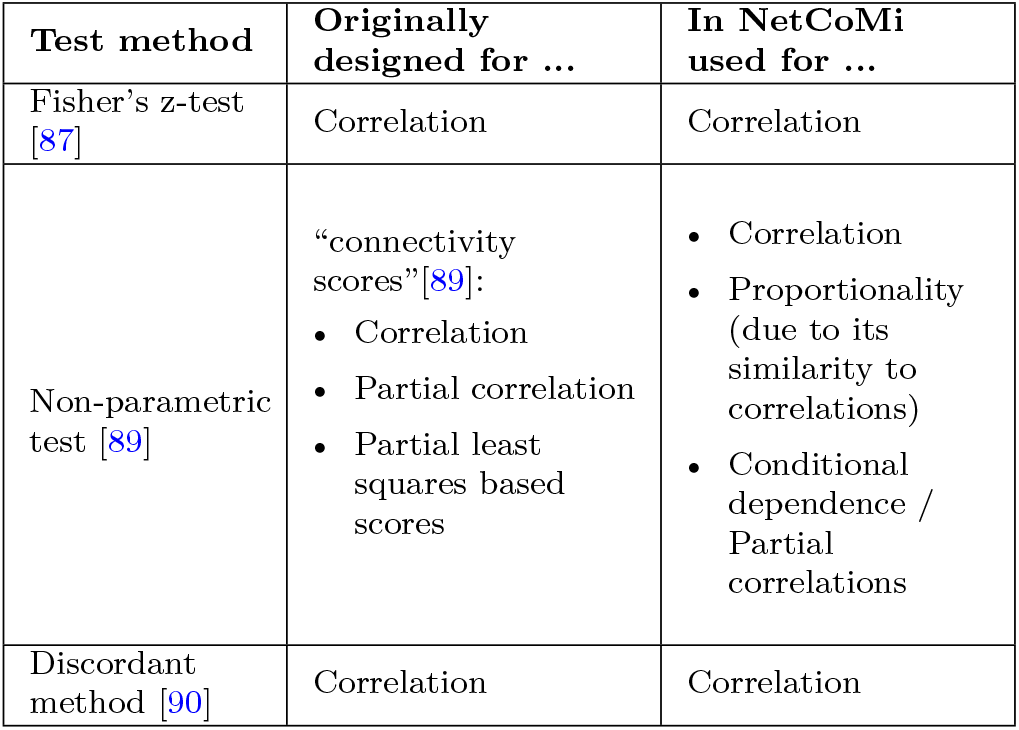
Test procedures for identifying differential associations available in `NetCoMi`. Fisher’s z-test and the Discordant method are based on correlations that are Fisher-transformed into z-values. These two methods are thus not suitable for the other association measures included in `NetCoMi`.

Fisher’s z-test [87] is a common method for comparing two correlation coefficients, assuming normally distributed z values (the transformed estimated correlations) and thus a correct specification of their variance. Novel approaches without normality assumption have been proposed [88], but are restricted to Pearson correlations. Therefore, we implemented a resampling-based procedure [89] as non-parametric alternative, which is applicable to association measures other than correlation. The Discordant method [90] as the third available method is also based on Fisher’s z values, but groups correlations with a similar magnitude and direction based on mixture models.

These approaches enable the construction of a differential network, where only differentially associated taxa are connected. More precisely, two taxa are connected if their association is either significantly different between the two groups (to a user-defined significance level) or identified as being different by the Discordant method.

## 4 Application of NetCoMi on a real data set

We use data from the GABRIEL Advanced Surveys (GABRIELA) [91] to illustrate the application of NetCoMi version 1.0.1 [92]. `GABRIELA` is a multi-center study, carried out in rural areas of southern Germany, Switzerland, Austria, and Poland, which provides new insights into the causes of the protective effect of exposure to farming environments for the development of asthma, hay fever, and atopy [91]. The study comprises the collection of biomaterial and environmental samples including mattress dust from children’s rooms and nasal swabs, for which 16S rRNA amplicon sequencing data are available. After some preprocessing steps (see Table S4), a total of *p* = 707 bacterial genera remain for mattress dust for a subset of *N* = 1022 subjects. The nasal data set consists of *p* = 467 genera for *N* = 1033 subjects.

The `NetCoMi` functions corresponding to the main steps of our proposed workflow (Fig. 1) are `netConstruct()` for network construction (Section 2.3), `netAnalyze()` for network characterization (Section 2.4), and `netCompare()` for network comparison (Section 3). An overview of the exported (and thus usable) `NetCoMi` functions together with their main arguments is given in Fig. 3. In the following, we use the SPRING method as a measure of partial correlation for constructing exemplary microbial association networks and Aitchison’s distance as a dissimilarity measure for constructing sample similarity networks. The construction of a differential network is described in Supplement 5.5.

**Fig. 3.**
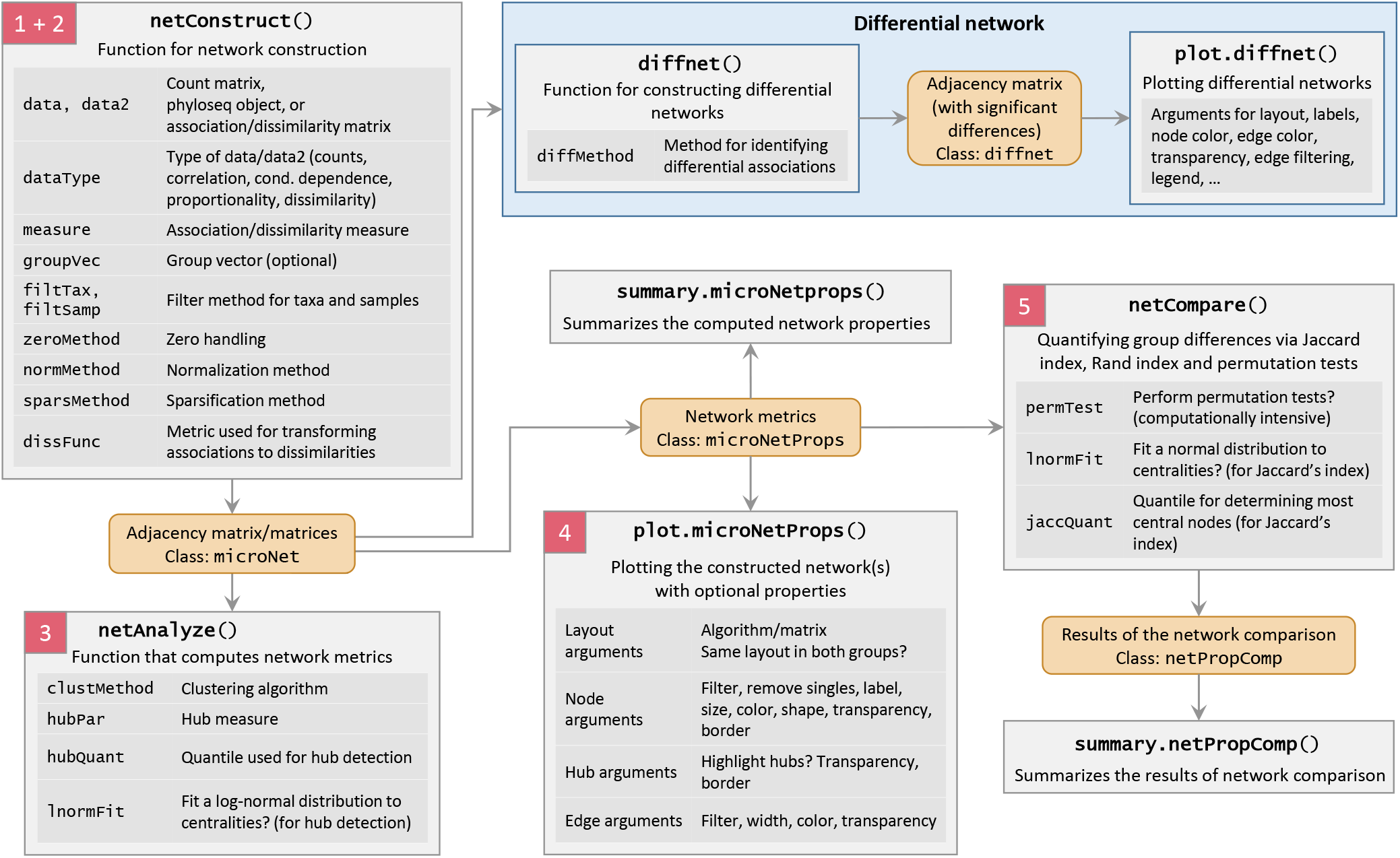
Main `NetCoMi` functions. For each function, its purpose together with its main arguments is shown. The objects returned from the respective functions are colored in orange. The steps (colored in red) correspond to the steps of the overall workflow shown in Fig. 1.

### 4.1. Constructing a single microbial network

In principle, the `netConstruct()` function allows the specification of any combination of methods for association measure, zero treatment, and normalization. However, since `NetCoMi` is mainly designed to handle compositional data, a warning is returned if a chosen combination is not compositionally aware.

To generate the network shown in Fig. 4A, we pass the combined count matrix containing dust samples from Ulm and Munich to `netConstruct()`. Filter parameters are set in such a way that only the 100 most frequent taxa are included in the analyses leading to a 1022 × 100 read count matrix for our data. Depending on the association measure, a method for zero handling, normalization and sparsification could be chosen (see Figure S1 and Table S1). Since these steps are already included in SPRING, they are skipped in our example. If associations have already been estimated in advance, the external association matrix can be passed to `netConstruct()` instead of a count matrix. In this mode, the user only needs to define a sparsification method and a dissimilarity function.

**Fig. 4.**
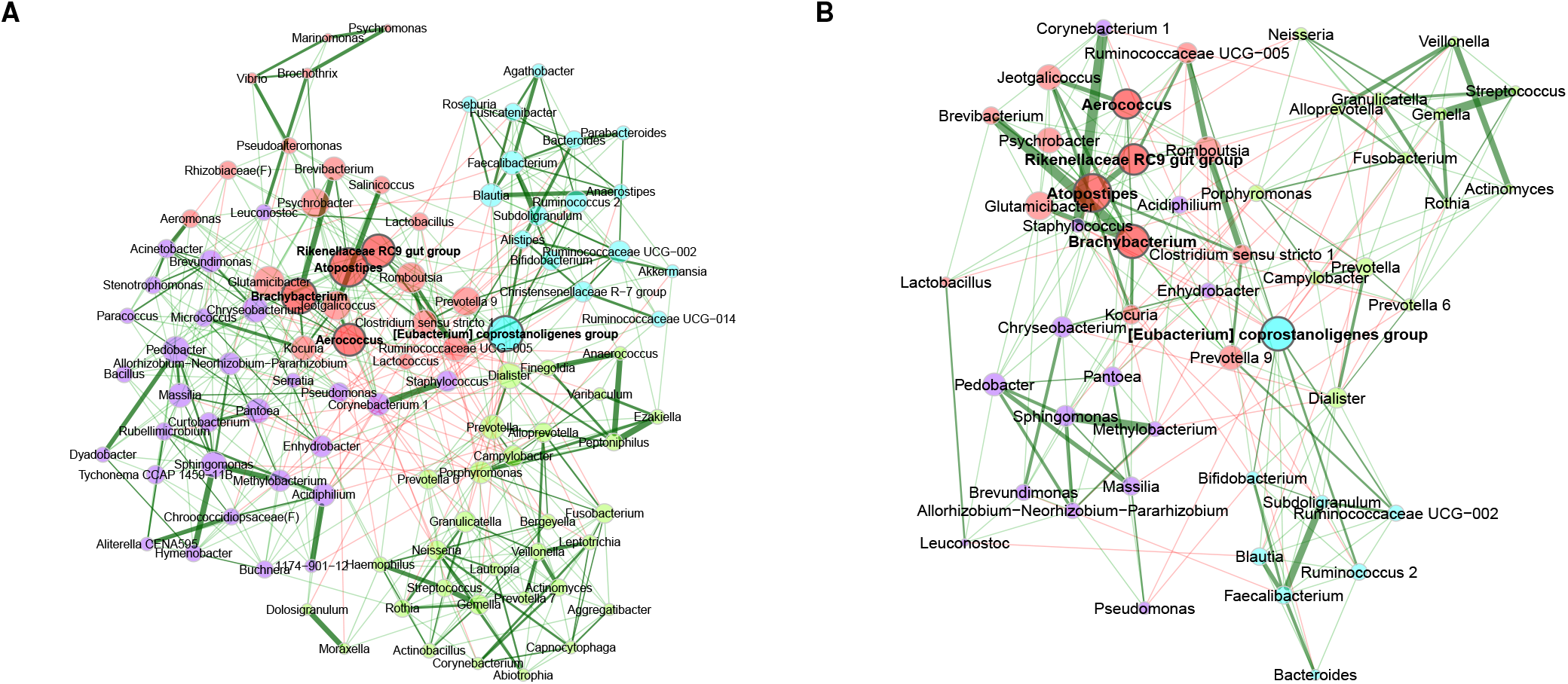
Bacterial associations for the combined data set with samples from Ulm and Munich. The SPRING method [39] is used as association measure. The estimated partial correlations are transformed to dissimilarities via the “signed” distance metric and the corresponding (non-negative) similarities are used as edge weights. Green edges correspond to positive estimated associations and red edges to negative ones. Eigenvector centrality is used for defining hubs (nodes with a centrality value above the empirical 95% quantile) and scaling node sizes. Hubs are highlighted by bold text and borders. Node colors represent clusters, which are determined using greedy modularity optimization. **A:** Complete network for the data set with 100 taxa and 1022 samples. Unconnected nodes are removed. **B:** Reduced network, where only the 50 nodes with the highest degree are shown. Centrality measures and clusters are adopted from the complete network.

Besides the available options for transforming the associations to dissimilarities (see Supplementary Figure S1), the function also accepts a user-defined dissimilarity function. The choice of dissimilarity function also influences the handling of negative associations. In Fig. 4, we use the “signed” distance metric, where strong negative correlations lead to a high dissimilarity (and thus to a low edge weight in the adjacency matrix). Figure S4 shows a network plot where the “unsigned” metric is used.

`netConstruct()` returns an object of class microNet, which can be directly passed to `netAnalyze()` to compute network properties (see Table S5). Applying the plot function to the output of `netAnalyze()` leads to a network visualization, as shown in Fig. 4A. In this plot, network characteristics are emphasized in different ways: Hubs are highlighted, clusters are marked by different node colors, and node sizes are scaled according to eigenvector centrality. Alternatively, node colors and shapes can represent features such as taxonomic rank. Node sizes can be scaled according to other centrality measures or absolute/relative abundances of the corresponding taxa. Node positions are defined using the Fruchterman-Reingold algorithm [93]. This algorithm provides a force-directed layout aimed at high readability of the network by placing pairs of nodes with a high absolute edge weight close together and those with low edge weight further apart.

The `plot()` method implemented in `NetCoMi` includes several options for selecting nodes or edges of interest to facilitate the readability of the network plot without influencing the calculated network measures. In Fig. 4B, for instance, we display the 50 bacteria with highest degree, facilitating network interpretability.

`NetCoMi` identified four clusters and the following five hub nodes: *Aerococcus*, *Atopostipes*, *Brachybacterium*, *Rikenellaceae RC9 gut group*, and *[Eubacterium] coprostanoligenes group*. The strongest positive association is between *Ezakiella* and *Peptoniphilus* with a partial correlation of 0.55 (in the green cluster in Fig. 4A). The strongest negative correlation is −0.09 between *Alistipes* (in the blue cluster) and *Lactobacillus* (in the red cluster).

### 4.2. Comparing networks between two study centers

For network comparison, the combined count matrix is again passed to `netConstruct()`, but this time with an additional binary vector assigning the samples to one of the two centers. This leads to the network plots shown in Fig. 5 (see Table S7 for the summary of network properties). The layout computed for the Munich network is used for both networks to facilitate the graphical comparison and making differences clearly visible. We observe only slight differences in the estimated associations. Furthermore, both networks show similar clustering and agree on three (out of five) hub nodes: *Atopostipes*, *Brachybacterium*, and *[Eubacterium] coprostanoligenes group*.

**Fig. 5.**
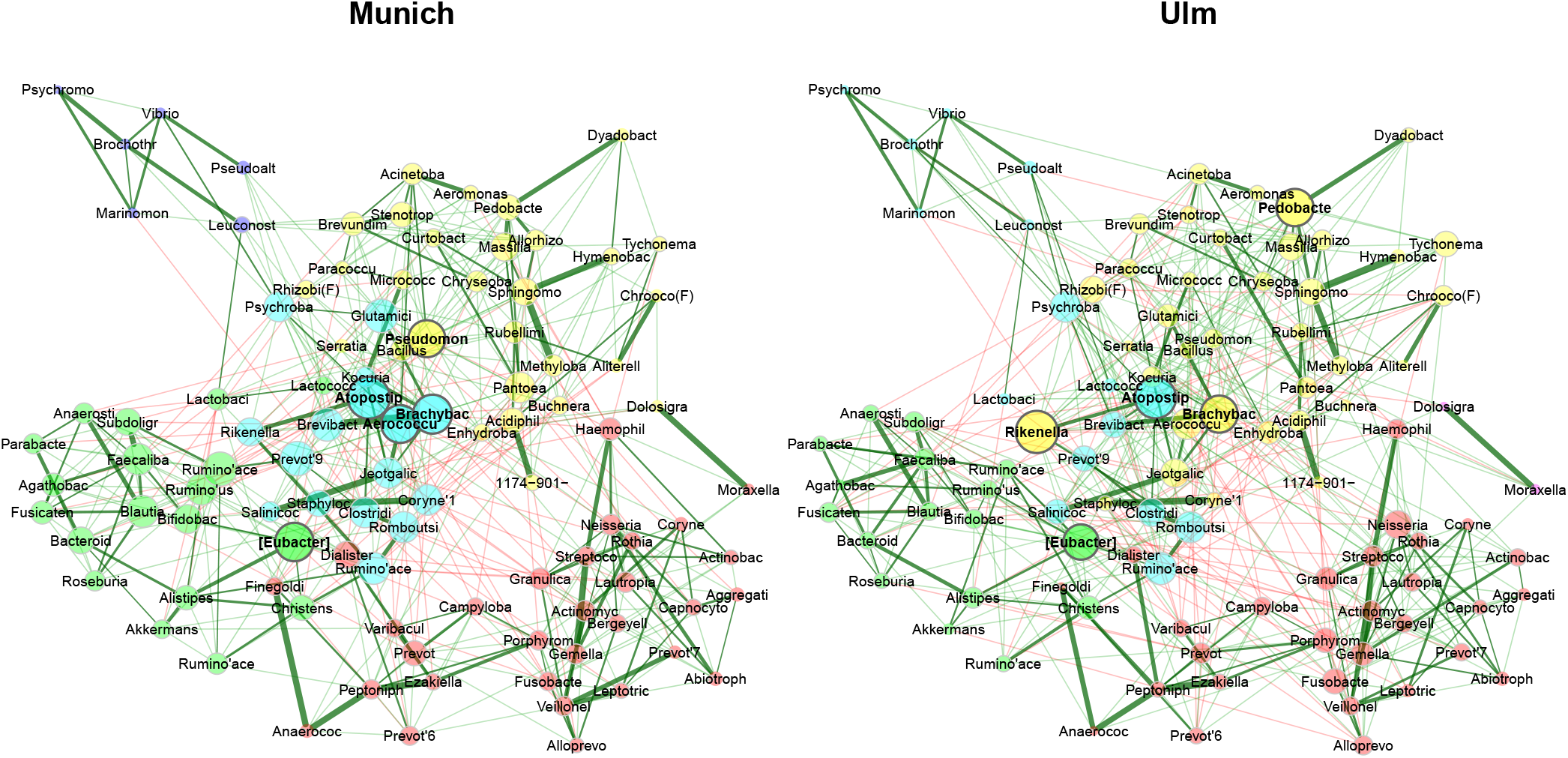
Comparison of bacterial associations in the mattress dust between the study centers Munich and Ulm. The SPRING method [39] is used as association measure. The estimated partial correlations are transformed to dissimilarities via the “signed” distance metric and the corresponding similarities are used as edge weights. Eigenvector centrality is used for defining hubs and scaling node sizes. Node colors represent clusters, which are determined using greedy modularity optimization. Clusters have the same color in both networks if they share at least 2 taxa. Green edges correspond to positive estimated associations and red edges to negative ones. The layout computed for the Munich network is used in both networks. Nodes that are unconnected in both groups are removed. Taxa names are abbreviated (see Table S9 for the original names).

The quantitative comparison is done by passing the R object returned from `netAnalyze()` to `netCompare()`. Comparisons of all global measures included in `NetCoMi` and the five genera with the highest absolute group difference for degree and eigenvector centrality, respectively, are given in Table 6. Supplementary Table S8 extends the output to the ten genera with the highest absolute group difference, and also includes betweenness and closeness centrality. For none of the four centrality measures, any significant differences are observed, confirming the descriptive analyses. *Pseudomonas*, *Pedobacter*, and *Rikenellaceae RC9 gut group* as hub taxa in only one of the networks show high differences in eigenvector centrality. However, the differences are not deemed significant (Table 6). Global network properties (upper part of Table 6) are also not significantly different for *α* = 5%.

**Table 6.** Results from testing global network metrics and centrality measures of the networks in Fig. 5 for group differences (via permutation tests using 1000 permutations). Shown are respectively the computed measure for Munich and Ulm, the absolute difference, and the p-value for testing the null hypothesis *H*_0_: *|*diff*|* = 0. For the centrality measures, p-values are adjusted for multiple testing using the adaptive Benjamini-Hochberg method [57], where the proportion of true *H*_0_ is determined according to Langaas et al. [58]. For degree and eigenvector centrality, the five genera with the highest absolute group difference are shown. The centralities are normalized to [0,1] as described in Table 4. Highly different eigenvector centralities (even if not significant) describe bacteria with highly different node sizes in the network plots in Fig. 5 such as *Neisseria*, which is a hub in Munich and much less important in Ulm.

|  | Munich | Ulm | abs. diff. | p-value |
| --- | --- | --- | --- | --- |
| Global network measures: |  |  |  |  |
| Average path length | 1.576 | 1.555 | 0.022 | 0.60440 |
| Clustering coefficient | 0.279 | 0.249 | 0.029 | 0.11189 |
| Modularity | 0.458 | 0.407 | 0.051 | 0.08192 |
| Vertex connectivity | 3 | 4 | 1 | 0.68232 |
| Edge connectivity | 3 | 4 | 1 | 0.68132 |
| Density | 0.103 | 0.100 | 0.003 | 0.55045 |
| Degree (weighted): |  |  |  |  |
| Corynebacterium 1 | 0.162 | 0.071 | 0.091 | 0.19905 |
| Rikenellaceae RC9 gut group | 0.101 | 0.192 | 0.091 | 0.82936 |
| Lactobacillus | 0.121 | 0.051 | 0.071 | 0.82936 |
| Pedobacter | 0.101 | 0.172 | 0.071 | 0.82936 |
| Pseudomonas | 0.202 | 0.131 | 0.071 | 0.99622 |
| Eigenvector centrality: |  |  |  |  |
| Pseudomonas | 0.852 | 0.417 | 0.435 | 0.57458 |
| Rikenellaceae RC9 gut group | 0.595 | 1.000 | 0.405 | 0.57458 |
| Ruminococcaceae UCG-002 | 0.767 | 0.379 | 0.389 | 0.57458 |
| Corynebacterium 1 | 0.670 | 0.290 | 0.379 | 0.57458 |
| Pedobacter | 0.526 | 0.873 | 0.346 | 0.83396 |
| Significance codes: ***: 0.001, **: 0.01, *: 0.05, .: 0.1 |  |  |  |  |

Table 7 summarizes Jaccard indices expressing the similarity of sets of most central nodes and the hub nodes among the two centers. They do not imply any group differences, which would be indicated by a small probability P(*J ≤ j*). Similarly, the adjusted Rand index (0.752, p-value = 0) indicates a high similarity of the two clusterings, which is also highlighted in Fig. 5.

**Table 7.** Jaccard index values corresponding to the networks shown in Fig. 5. Index values *j* express the similarity of the sets of most central nodes and also of the sets of hub taxa between the two networks. “Most central” nodes are those with a centrality value above the empirical 75% quantile. Jaccard’s index is 0 if the sets are completely different and 1 for exactly equal sets. *P* (*J ≤ j*) is the probability that Jaccard’s index takes a value less than or equal to the calculated index *j* for the present total number of taxa in both sets (*P* (*J ≥ j*) is defined analogously).

| | $j$ | $P(J \leq j)$ | $P(J \geq j)$ |
| --- | --- | --- | --- |
| degree | 0.323 | 0.53375 | 0.61683 |
| betweenness centr. | 0.389 | 0.81277 | 0.29332 |
| closeness centr. | 0.471 | 0.96735 | 0.06729 |
| eigenvec. centr. | 0.429 | 0.91326 | 0.15482 |
| hub taxa | 0.429 | 0.82670 | 0.42936 |
| Significance codes: ***: 0.001, **: 0.01, *: 0.05, .: 0.1 |  |  |  |

### 4.3. Sample similarity networks

We next consider sample similarity networks from mattress dust and nasal swabs of the same subjects (*N* = 980). The process of constructing and comparing sample similarity networks is analogous to the one for association networks. Figure 6 shows the two networks using Aitchison’s distance.

**Fig. 6.**
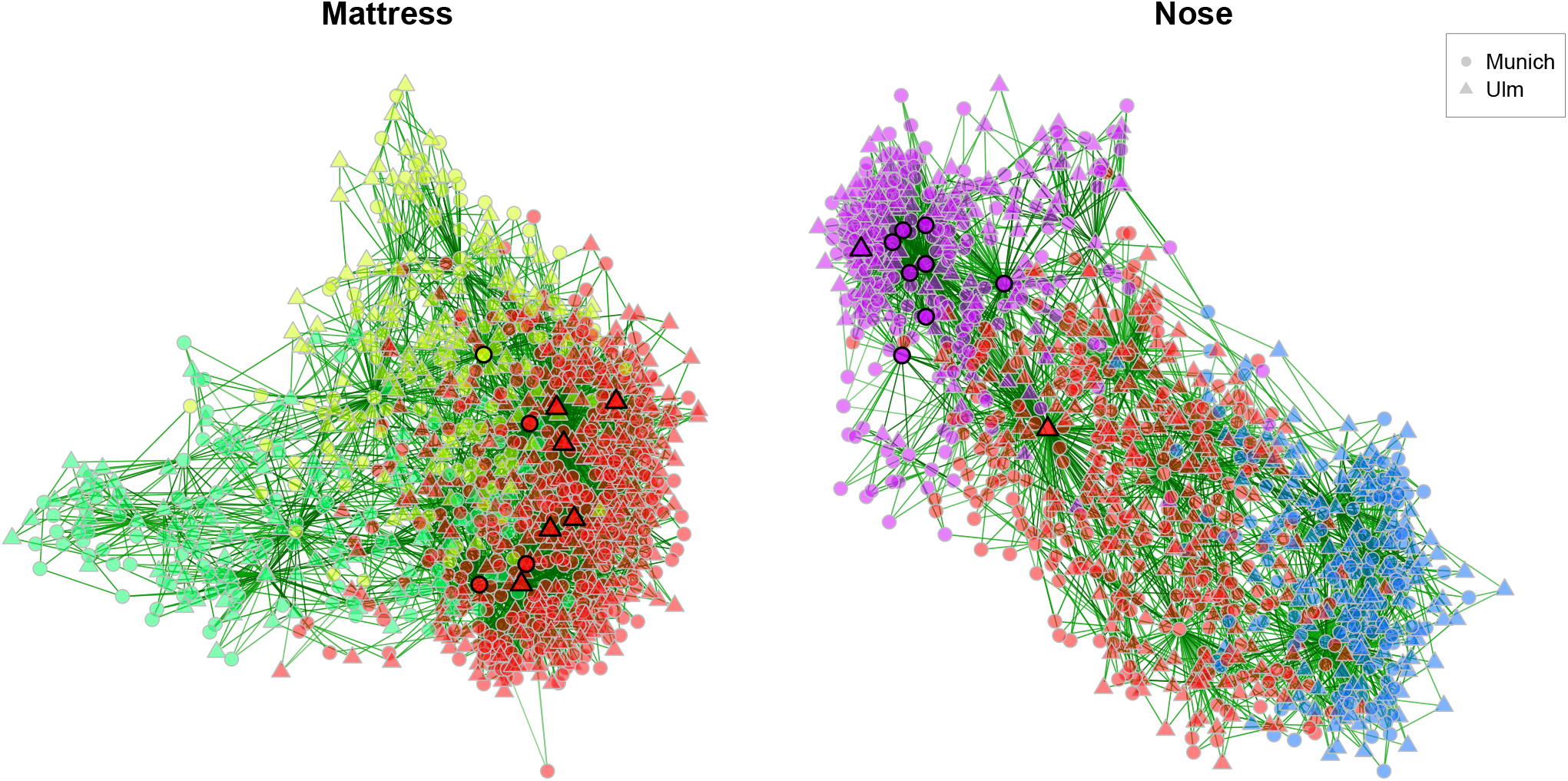
Comparing dissimilarity networks based on Aitchison’s distance [80] (see Supplementary Table S3) between mattress dust and nasal swabs for the same set of subjects (nodes). Only samples and taxa with at least 1000 reads, respectively, are included leading to *p*_1_=707 genera in the Mattress group, *p*_2_=184 genera in the Nose group, and *n*=980 samples in both groups. Counts are normalized to fractions and – since zeros must be replaced for the clr transformation – “multiplicative imputation” (see Table 3 in the main text) is used for zero handling. The dissimilarity matrix is scaled to [0,1] and sparsified using the k-nearest neighbor method (*k*=3 for both networks). Node colors represent clusters, identified using hierarchical clustering with average linkage. A cluster has the same color in both networks if they have at least 100 nodes in common (the minimum cluster size among both groups is 560). Hubs (highlighted by bold borders) are nodes with an eigenvector centrality larger than the 99% quantile of the empirical quantile of eigenvector centralities. Edge thickness corresponds to similarity values (calculated by 1 − *distance*). Nodes are placed further together, the more similar their bacterial composition is. Whether a sample has been collected in Munich or Ulm is marked by node shapes. Unconnected nodes are removed.

`NetCoMi`’s quantitative network analysis (Table S10 to S13) reveals strong differences between these networks. The sets of most central nodes are significantly different for all four centrality measures (shown by Jaccard’s index). Hub nodes are also completely different (Jaccard index of zero). We also observe several significantly different global network properties (see Table S12). Furthermore, the node’s degree, betweenness, and closeness centrality values differ significantly between the groups for several subjects (Table S13), implying that a single subject plays a different role dependent on the investigated microbial habitat.

Clustering analysis broadly identifies three sample groups for mattress dust and nasal swab samples, respectively. In Fig. 6, we highlight the partial overlap between the two clustering solutions via color coding. Clusters that have at least 100 nodes in common are plotted by matching colors. This reveals that the red cluster, seen in both networks, shares similar samples across the two habitats.

`NetCoMi`’s plotting functionality also allows to draw nodes in different shapes, corresponding to additional categorical covariate information about the samples. For instance, in Fig. 6, the two node shapes correspond to the two different study centers (Ulm and Munich). This feature could potentially highlight confounding of groups of samples and available covariates. For our example here, however, we observe no noticeable pattern in the clusters with respect to study center.

## 5 Discussion

### 5.1. Why use NetCoMi?

With `NetCoMi` we offer an easy-to-use and versatile, integrative `R` package for the construction, analysis, and comparison of microbial networks derived from amplicon sequencing data. Our package provides a wide variety of compositionally aware association measures, including SparCC [5], proportionality [38], SPIEC-EASI [47], and SPRING [39, 7]. The latter method also enables the analysis of recent quantitative microbiome data sets when both amplicon and quantitative cell count or spike-in control data are available. `NetCoMi` also incorporates standard association measures, thus widening the scope of the package beyond applications to compositional data, and it connects to the popular `WGCNA` package [35], enabling principled *soft-thresholding* of correlations and dissimilarity transformations based on topological overlap. The package includes a dedicated list of methods for handling excess zeros in the count matrix and for data normalization in order to account for the special characteristics of the underlying sequencing data prior to association estimation.

A unique feature of the `NetCoMi` framework is its ability to perform differential network and differential association analysis in a statistically principled fashion. Differential network analysis allows not only to uncover the global role of a taxon in the overall network structure but also its changing influence under varying conditions. Differential association analysis [14], on the other hand, can directly assess which associations significantly change across conditions, providing concrete hypotheses for follow-up biological perturbation experiments.

Similar to `phyloseq`’s [94] `plot_net` function, `NetCoMi` also enables network representation and comparison of the amplicon data samples themselves, using popular sample dissimilarity or distance measures, such as the Bray-Curtis dissimilarity and the Aitchison distance. Network analysis of the resulting sample-to-sample or subject-to-subject networks can give insights into the heterogeneity of the collected data. For instance, identified *hub subjects* are subjects with “representative” bacterial compositions that may comprise archetypical microbial patterns in the studied population. Sample similarity network analysis thus extends standard sample ordination or cluster analysis, making fine-grain structures of the available microbial sample collection visible.

While methods and tools for the individual analysis steps, such as biological network estimation (see, e.g. [95, 96] for recent contributions) and (differential) biological network analysis [17, 16, 35, 97, 94, 15] are available, `NetCoMi` offers a unique and modular `R` software framework that integrates the complete process of estimating, analyzing, and comparing *microbial* networks. From an end user’s perspective with a specific microbiome dataset and scientific question in mind, `NetCoMi` will thus facilitate both faster development and reproducibility of the microbial network analysis workflow.

### 5.2. Which method to choose?

Even though the modular design of `NetCoMi` allows the user to perform a wide variety of computational workflows, going from the primary data all the way to potentially significant network features, every step of the analysis still warrants careful scientific consideration. For instance, the choice of an association or dissimilarity measure will likely affect all further steps of network analysis and comparison. However, there is no general consensus in the community about the “right” way to estimate and analyze microbial networks. This is reflected in the heterogeneity of recent simulation studies to assess and compare the performance of compositionally aware as well as traditional association measures. To put some of these studies into context, we give a selection of simulation studies examining the association methods used in `NetCoMi` in Supplementary Table S14.

Common shortcomings in current simulation studies include the lack of a universal standard (i) to generate realistic synthetic microbial data with a prescribed ground truth, (ii) to perform comparable model selection, and (iii) to report generalizable performance metrics. Comparative studies often concentrate on the performance in edge recovery, for instance, via precision-recall curves [39, 98, 6, 99] or distances between true and estimated associations [5, 36, 39, 6]. However, networks derived from penalized marginal correlations (such as SparCC) and partial correlation (such as SPRING) are statistically difficult to compare, thus requiring special modifications which, in turn, limits cross-study comparisons (see, e.g. Yoon et al. [39] where SparCC correlations are transformed to SparCC partial correlations). Moreover, the sole focus on edge recovery may obfuscate other aspects of correct model recovery, including the shape of degree distributions [6] or detecting hub nodes. For instance, methods with a similar edge recovery precision greatly vary regarding their ability to determine hub nodes [100]. Thus, if the correct detection of hub nodes is of major interest to the user, present comparative microbial network studies will give little guidance.

Finally, we posit that simulation studies accompanying articles that introduce new methods might also be inherently biased [101, 102]. Neutral comparison studies, i.e., studies that are independent of any new method development [101, 102] are rare in our context or do not include recently published methods [98]. The main impediment regarding neutral comparison studies is, however, the fact that, to date, no large-scale ground truth network of real biological microbial interactions is available. Such biological gold-standard networks, as available in other contexts (e.g. gene-gene or transcription-factor-gene interactions), would greatly facilitate future comparative studies.

In the absence of a “best method” for microbial network inference and analysis, `NetCoMi` is intended to give researchers the possibility to apply a consistent and reproducible analysis workflow on their data. Ideally, the selection of the workflow building blocks should be set up once and, independent of any hypothesis about the data, thus avoiding the fallacy of starting “fishing” for results that best suit a previously formulated hypothesis. `NetCoMi` can, however, serve as an ideal tool for principled sensitivity analysis of the inferred results, for instance, by assessing how different normalization and zero handling methods affect the estimated networks, their structural properties, and their comparison. Finally, we envision `NetCoMi` to provide a useful framework for future simulation studies that evaluate and compare the performance of different association measures and network inference tools in a reproducible fashion.

### 5.3. Current limitations and future developments

The current version of `NetCoMi` is designed to model networks from a single domain of life, e.g. bacteria, fungi, or viruses. However, microbes from different domains of life often share the same habitat and likely influence each other [103]. Joint cross-domain network inference already revealed considerable alterations of the overall network structure and network features, compared to their single-domain counterparts [104]. Extending `NetCoMi` to cross-domain network analysis is thus an important future development goal. Likewise, environmental factors, such as chemical gradients and temperature, as well as batch effects are known to influence microbial abundances and composition and thus bias network estimation [100, 105, 106]. In the current `NetCoMi` version, we assume that the user has already corrected the microbiome data for these latent influences. However, several inference methods can directly incorporate known [107, 108] or unknown latent factors [105] into network learning. Including or connecting these approaches with `NetCoMi` will likely increase the robustness and generalizability of future workflows.

A core feature of `NetCoMi` is the use of statistical tests at various stages of the computational workflow. For instance, statistical tests can be employed for edge selection in network sparsification. Since statistical power depends on sample size, the sparsified structure of a network will likely depend on the number of available samples. A comprehensive understanding between number of samples, sparsification, and network structure is currently elusive. `NetCoMi` relies on permutation tests for several statistical tests. The lower limit of p-values arising from permutation tests is directly related to the number of available permutations (Section 4). This already required proper adjustment of the calculation for small numbers [109, 110]. For extended simulation studies, `NetCoMi`’s dependence on permutation tests may prove to be computer intensive. Thus, integration of less demanding alternatives to permutation tests would represent a welcome feature in `NetCoMi`.

Finally, despite incorporating a comprehensive list of methods in our `R` package, we do not claim completeness. This can hardly be achieved in the vibrant field of microbiome research where new methods are constantly developed. We alleviated this shortcoming via `NetCoMi`’s modular structure which allows certain parts of our workflow to be combined with external methods. For instance, users can input a user-defined association or dissimilarity matrix rather than a data matrix, and then proceed with `NetCoMi`’s standardized network analysis modules.

In summary, we believe that `NetCoMi` is a useful addition to the modern microbiome data analysis toolbox, enabling rapid and reproducible microbial network estimation and comparison and ideally leading to robust hypotheses about the role of microbes in health and disease [111].

##### Key Points

- Current high-throughput amplicon sequencing count data carry only relative or compositional information, thus requiring dedicated statistical analysis methods.
- `NetCoMi` is a comprehensive `R` package that implements the complete workflow of constructing, analyzing, and comparing microbial association networks.
- `NetCoMi` integrates an extensive list of methods that take into account the special characteristics of amplicon data, including methods for zero count handling, normalization, and association estimation.
- The package also offers functionality for constructing sample similarity networks as well as differential networks including appropriate methods for identifying differentially associated taxa.

## Supporting information

R Script

## Acknowledgments

We gratefully thank Hanna Danielewicz (Wroclaw Medical University), Caroline Roduit (University Children’s Hospital Zurich), Dick Heederik (Utrecht University), and Jon Genuneit (Ulm University and Leipzig University) who were responsible for design and implementation of the GABRIELA study as well as Rob Knight and Georg Loss (both UC San Diego) who created the 16S rRNA data used in our examples. We would like to thank Alethea Charlton for language editing and her helpful comments. We also thank Anastasiia Holovchak for testing the `R` package and editing its documentation.

## Data availability

The data underlying this article will be shared on reasonable request to the corresponding author upon approval by the involved institutional review boards.

## Author information

**Stefanie Peschel** is a PhD student at the Institute for Asthma and Allergy Prevention, Helmholtz Zentrum München (Munich, Germany) with background in statistics. Her current research focuses on understanding the role of the human microbiome for the development of asthma and allergies using network analysis methods.

**Christian L. Müller** is group leader at the Institute of Computational Biology, Helmholtz Zentrum München, professor at the Department of Statistics, LMU München, and researcher at the Center for Computational Mathematics, Flatiron Institute, New York. His research interests include computational statistics, optimization, and microbiome data analysis.

**Erika von Mutius**, MD MSc, Professor of Pediatric Allergology, is head of the Asthma and Allergy Department at the Dr. von Hauner Children’s Hospital in Munich and leading the Institute for Asthma and Allergy Prevention at Helmholtz Zentrum Munich. Her research group is focusing on the determinants of childhood asthma and allergies.

**Anne-Laure Boulesteix** is an associate professor at the Institute for Medical Information Processing, Biometry and Epidemiology, University of Munich (Germany). Her activities mainly focus on computational statistics, biostatistics and research on research methodology.

**Martin Depner** is a postdoctoral research fellow at the Institute for Asthma and Allergy Prevention, Helmholtz Zentrum München (Munich, Germany) with background in medical biometry. His current research focuses on studies of the microbiome (16S rRNA) and the protective effects of growing up on a farm on asthma and allergies.

