## Supplementary Material for "NetCoMi: Network Construction and Comparison for Microbiome Data in R"

**6 Existing simulation studies comparing association and dissimilarity measures** **25**

### 1 Further details on association measures

#### 1.1 Correlation measures

##### 1.1.1 SparCC

SparCC (Sparse Correlations for Compositional data) (Friedman and Alm, 2012) estimates Pearson correlations of the true unobserved abundances via an approximative approach based on the variance of log-ratios $t_{ij} = \text{Var}\left(\log\frac{x_i}{x_j}\right)$. The approximation is valid under the assumptions that the number of taxa is large and that they are sparsely correlated.

Since log-ratios cannot be calculated if the count matrix contains any zeros, they need to be handled suitably. Friedman and Alm (2012) propose a Bayesian approach, where the true (unobserved) fractions, given the observed counts, are assumed to be Dirichlet distributed. All observations are replaced by random samples from that posteriori. The sparsity assumption is reinforced by iterating the process of correlation estimation, where the taxa pair with the highest estimated correlation is excluded in each step as long as their correlations exceed a predefined threshold. Then, the process of estimating fractions and computing correlations is repeated. To account for variation in the sampling process, the whole approach is repeated several times. The authors of SparCC suggest at least 100 repetitions, which is used as default value in `NetCoMi`.

We adopted some code from the `r-sparcc` package (Filosi, 2017) to implement a parallelized version of the SparCC algorithm so that the repetitions of the basic procedure are executed parallel using multiple CPU cores. In `NetCoMi`, the basic algorithm is implemented in C++ (using the R package `Rcpp`) to further reduce run time.

##### 1.1.2 CCLasso

CCLasso (Correlation inference for Compositional data through Lasso) (Fang et al., 2015) is another approach for inferring the latent correlations between microorganisms. In CCLasso, log-ratios are used to overcome compositional effects as well. Furthermore, a sparsity assumption is necessary to infer the covariance matrix, and thus the correlation matrix, from the observed compositional data. The basis of CCLasso is an algorithm for finding a positive definite correlation matrix with elements in [-1,1] by minimizing a loss function plus an $l_1$-penalty. The loss function considers errors resulting from estimating the covariance matrix from the observed compositional data instead of the true abundances and the penalty incorporates the sparsity assumption. The R code for CCLasso published on GitHub (Fang, 2016) has been adopted to `NetCoMi`.

##### 1.1.3 CCREPE

CCREPE (Compositionality Corrected by REnormalization and PErmutation), originally called "ReBoot method", has been proposed by Faust et al. (2012). CCREPE operates on the observed relative abundances. CCREPE uses a procedure to test for statistical significance of the estimated correlations that takes into account spurious correlations caused by compositional effects.

The idea is that the null distribution constructed via permutation should incorporate that a part of the observed correlation is caused by compositionality alone. To construct an appropriate permutation distribution, the relative abundances are permuted and re-normalized so that they sum up to one within a sample again. The correlations between organisms are calculated for the permuted data, and this process is repeated to obtain a null distribution which should represent correlations caused by compositionality alone. Then, a bootstrap confidence interval is constructed which is located around the "observed“ correlation. Finally, a z-test is used to test for differences in the mean of both distributions. A significant result implies that the estimated correlation for the observed data is significantly different from the correlation induced by compositionality. For the CCREPE approach, a zero treatment is possible but not necessarily needed.

As stated in Faust et al. (2012), relative data do not contain all information about the true absolute abundances, which is why it is generally not possible to fully distinguish between spurious correlations due to compositionality from original correlation Faust et al. (2012).

##### 1.2 Measures of conditional independence

###### 1.2.1 SPIEC-EASI

A basic assumption of SPIEC-EASI (Sparse Inverse Covariance Estimation for Ecological Association Inference) (Kurtz et al., 2015) is that the covariance matrix for *clr*-transformed data is approximately equal to the ”true“ basis covariance matrix if the number of taxa $p$ is large. After *clr*-transformation, an undirected weighted graph is estimated using one of two provided methods: neighborhood selection (MB) or sparse inverse covariance selection (GL). In the MB approach, for every single node, a penalized regression model is solved. The GL method, in contrast, seeks to reconstruct the whole graph by solving a global convex optimization problem. In the $p > n$ setting, both methods assume the inverse covariance to be sparse.

The neighborhood selection approach leads to node-wise partial correlations for each node that after symmetrization, can serve as weighted adjacency matrix. The GL approach results in a positive definite inverse covariance matrix (precision matrix) $\Sigma^{-1}$, whose negative entries are proportional to partial correlations. The non-zero entries of the matrix form the adjacency matrix. Both methods include a tuning parameter $\lambda$ controlling the sparsity of the estimates. Model selection via the “Stability Approach to Regularization Selection (StARS)”. Here, random subsamples of the data are taken repeatedly, from which graphs are estimated across the entire regularization path. From the resulting collection of graphs, an overall edge variability statistic is computed across the $\lambda$-path. The StaRS-selected $\lambda$ is the smallest $\lambda$ at which a user-defined stability threshold is reached.

###### 1.2.2 SPRING

The SPRING (Semi-Parametric Rank-based approach for INference in Graphical model) approach (Yoon et al., 2019) follows a similar workflow as SPIEC-EASI but is applicable both to quantitative and relative abundance data. In SPRING, the correlation matrix is estimated via a semi-parametric rank-based correlation estimator. The data are assumed to follow a truncated Gaussian copula model (Yoon et al., 2018), which extends the original Gaussian copula model (Liu et al., 2012) in order to handle data with excess zeros. The latent correlation matrix is estimated using Kendall’s $\tau$ and element-wise univariate optimization of an inverse copula function.

Sparse conditional dependencies are then inferred using the neighborhood selection approach using the latent correlation estimate. Analogous to SPIEC-EASI, model selection is performed via StARS. When applied to compositional data, SPRING introduces a modified $clr$ transformation ($mclr$) that does not require pseudocounts. The $mclr$ transform calculates the geometric mean from positive proportions only, computes the $clr$ for all positive parts with respect to this mean, and applies a positive shift to all non-zero components to ensure strict positivity. Zero proportions remain zero in the $mclr$-transformed vector.

###### 1.2.3 gCoda

The basic assumption of gCoda (Fang et al., 2017) is that the unobserved absolute read counts $y$ follow a multivariate normal distribution with mean $\mu$ and covariance matrix $\Sigma$, which describes the dependency structures between variables and thus is the term of interest. From $\log(y) \sim N_p(\mu, \Sigma)$ follows that the observed relative counts are logistic normal distributed. In gCoda, also a sparsity assumption is needed to determine the covariance matrix. To satisfy this assumption –analogous to CCLasso– a $l_1$ penalty is used, which is added to the negative log-Likelihood. The resulting optimization problem is solved via a Majorization-Minimization algorithm to estimate the inverse covariance matrix.

#### 2 Further details on network construction

##### 2.1 Detailed workflow for determining the adjacency matrix from associations

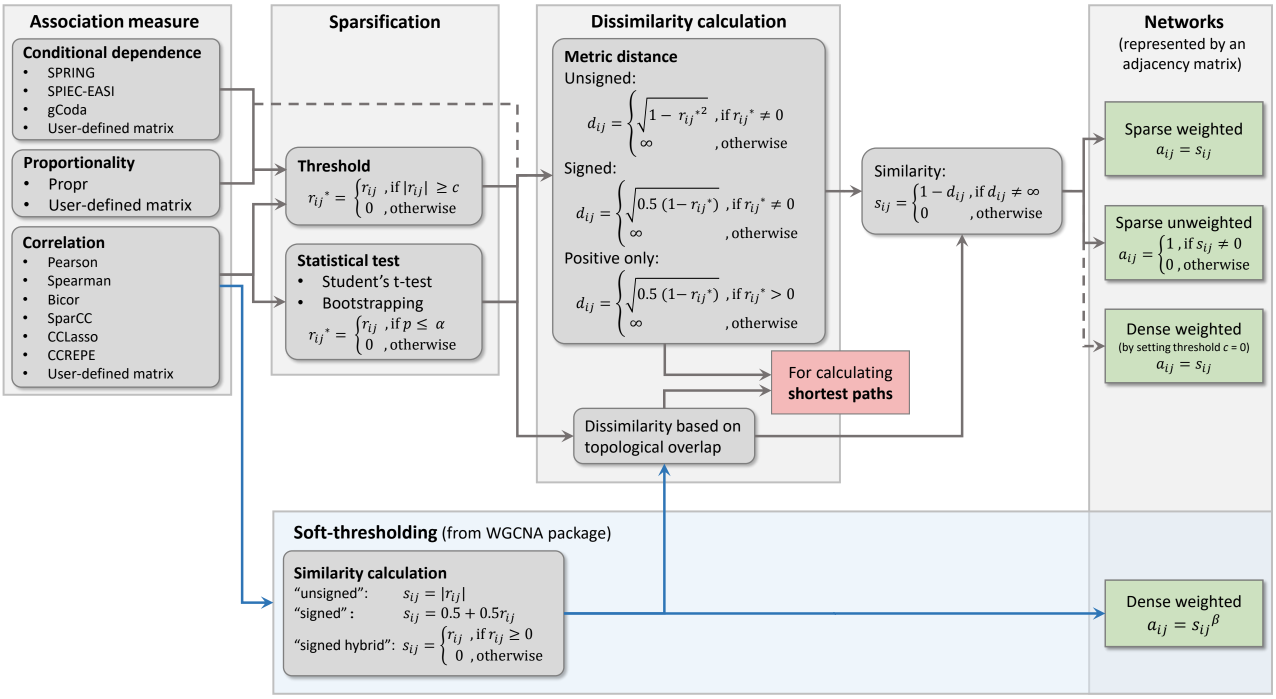

Figure S1: Approaches for constructing the adjacency matrix with entries $a_{ij}$ from estimated associations $r_{ij}$ that are available in NetCoMi. Depending on the association measure, the user can choose between different sparsification methods. The associations are then transformed into dissimilarities $d_{ij}$, which are used for network properties based on shortest paths. The dissimilarity between nodes with $r^*_{ij} = 0$ is set to infinity so that a direct path exists only between nodes with an association different from zero. The dissimilarities are, in turn, transformed into similarities via $s_{ij} = 1 - d_{ij}$. Similarities are used for similarity-based network properties as well as edge weights in the network plot. For correlations, in addition to a threshold and statistical testing, the soft-thresholding approach from the WGCNA package (Langfelder and Horvath, 2008) is available (blue path). The WGCNA package offers three possibilities for transforming correlation into similarity: "unsigned", "signed", and "signed hybrid", which are also available in NetCoMi. Following Langfelder and Horvath (2008), these similarities are used as edge weights and for similarity-based network measures, whereas for shortest paths a dissimilarity measure based on the topological overlap matrix (TOM) is used. The TOM-based dissimilarity has generally been made available as dissimilarity transformation in NetCoMi.

##### 2.2 The WGCNA approach

###### 2.2.1 Soft-thresholding

WGCNA has originally been proposed by Zhang and Horvath (2005) as "Weighted Gene Co-expression Network Analysis", which later led to the R package WGCNA ("weighted correlation network analysis") (Langfelder and Horvath, 2008). In this approach, a "soft thresholding" method is used for network construction as an alternative to common edge selection methods. The Zhang and Horvath (2005) argue that "hard" thresholding, such as hypothesis testing or a cutoff value, goes along with a loss of information. This is plausible because only selected edges are part of the network, and thus, are included in further analyses such as cluster building or calculating centrality measures. The authors propose to raise the estimated correlation to the power of a predefined value $\beta$ to determine the adjacencies. For $\beta > 1$, the adjacency increases not linearly anymore. Instead, small correlation values are pushed towards zero so that they become less important in the network. An overview, how connection strength changes for different powers $\beta$ is given in Mason et al. (2009).

###### 2.2.2 Main steps of the WGCNA approach

The idea of the WGCNA approach (Zhang and Horvath, 2005) is as follows: The estimated correlations between every two variables $i$ and $j$ are transformed into similarities $s_{ij}$. The entries of the adjacency matrix are then defined as $a_{ij} = s^{\beta}_{ij}$. As the next step, dissimilarity values based on the topological overlap measure (TOM) are determined from the estimated similarities. These dissimilarities are the basis for determining clusters or modules via a hierarchical clustering algorithm.

Thus, in contrast to our proposed workflow (where the correlations are transformed to distances first, which are then, in turn, transformed to similarities), in WGCNA the similarity values are directly calculated from the correlations.

The `WGCNA` package (Langfelder and Horvath, 2008) offers different options for transforming correlations to similarities leading to three network types, which handle negative correlations in different ways. The proposed network types are: "unsigned","signed", and "signed hybrid", with the following corresponding transformations:

$$s_{ij}^{\text{unsigned}} = |r_{ij}|,$$

$$s_{ij}^{\text{signed}} = 0.5 + 0.5\, r_{ij},$$

$$s_{ij}^{\text{signed hybrid}} = \begin{cases} r_{ij}, & \text{for } r_{ij} > 0 \\ 0, & \text{otherwise.} \end{cases}$$

Thus, the estimated correlations, which are located in [-1,1] are transformed into similarities ranging from 0 to 1.

The power $\beta$ is determined based on a scale-free criterion. The idea is that the degree distribution $P(k)$, which is defined as the fraction of vertices in a network with $k$ adjacent nodes, follows a power-law distribution: $P(k) \sim k^{-\gamma}$ for most biological networks. A linear regression model is used to assess how well $P(k)$ fits a power-law distribution. Networks leading to an $R^2 > 0.8$ are assumed to be approximately scale-free (Zhang and Horvath, 2005). To choose an appropriate power $\beta$, the authors suggest generating the adjacency matrix for different $\beta$ values and choose the smallest power for which the scale-free topology is fulfilled. This can be done in R via the `pickSoftThreshold` function (Langfelder and Horvath, 2008). Alternatively, a user-defined power parameter is accepted as input.

We adopted the aforementioned approaches of the soft-thresholding approach from the `WGCNA` package to `NetCoMi` via own `R` code. Thus, the user of our package can choose between one of the options "signed", "unsigned" and "signed hybrid" to generate similarities, which are transformed to adjacencies via $s_{ij}^{\beta}$. The power parameter can either be defined in advance or computed automatically. In the latter case, the `pickSoftThreshold()` function is used. Following the WGCNA approach Zhang and Horvath (2005), TOM-based dissimilarities are used as the basis for clustering, but also for calculating shortest paths.

##### 2.3 Notes on normalization and zero handling

Table S1: Overview of association measures implemented in `NetCoMi`. For each measure is shown, whether it is suitable for read count data without transforming the data in advance, whether a zero treatment is necessary and which normalization is accepted in `NetCoMi`. The function for network construction returns a warning when the chosen combination of association measure and normalization method may lead to compositional effects.

| Measure | | Compositionally aware? | Zero treatment | Normalization |
|---|---|---|---|---|
| Pearson coef. | Correlation | no | not needed | any |
| Spearman coef. | Correlation | no | not needed | any |
| Biweight midcorr. | Correlation | no | not needed | any |
| SparCC | Correlation | yes | included | none |
| CCLasso | Correlation | yes | needed | fractions |
| CCREPE | Correlation | yes | not needed | fractions |
| ρ | Proportionality | yes | needed | none |
| SPRING | Conditional dependence | yes | included | none |
| SPIEC-EASI | Conditional dependence | yes | included | none |
| gCoda | Conditional dependence | yes | needed | fractions |
| Euclidean distance | Dissimilarity / distance | no | not needed | any |
| Bray-Curtis dissim. | Dissimilarity / distance | no | not needed | any |
| Kullback-Leibler Divergence (KLD) | Dissimilarity / distance | no | needed | any |
| Jeffrey divergence | Dissimilarity / distance | no | needed | any |
| Jensen-Shannon diverg. | Dissimilarity / distance | no | needed | any |
| Compositional KLD | Dissimilarity / distance | yes | needed | fractions |
| Aitchison distance | Dissimilarity / distance | yes | needed | fractions |

##### 2.4 Methods for sample and taxa filtering implemented in NetCoMi

Table S2: Methods for preprocessing the count matrix by filtering samples and taxa implemented in `NetCoMi`. Except for `highestFreq` and `highestVar`, several methods can be combined. The parameter $x$ is user-defined.

| Argument name | Description |
|---|---|
| Sample filtering: | |
| `totalReads` | Keep samples with a total number of reads of at least $x$. |
| `numbTaxa` | Keep samples for which at least $x$ taxa are observed. |
| `highestFreq` | Keep the $x$ samples with highest frequency. |
| Taxa filtering: | |
| `totalReads` | Keep taxa with a total number of reads of at least $x$. |
| `relFreq` | Keep taxa whose number of reads is at least $x$% of the total number of reads. |
| `numbSamp` | Keep taxa observed in at least $x$ samples. |
| `highestVar` | Keep the $x$ taxa with highest variance. |
| `highestFreq` | Keep the $x$ taxa with highest frequency. |

#### 2.5 Sample similarity networks

NetCoMi provides the possibility to use a dissimilarity measure for network construction instead of association measures. The available measures are shown in Table S3. Using one of these measures leads to a network where nodes are subjects/samples instead of taxa. Aitchison's distance, the compositional Kullback-Leibler divergence (KLD) as well as the Bray-Curtis index are developed for quantifying differences in biological communities. In particular, the Aitchison distance and compositional KLD are designed for compositional data, but also the Bray-Curtis dissimilarity is appropriate for the application on compositional data (Weiss et al., 2016).

The complete workflow for constructing a dissimilarity-based network is shown in Figure S2. Data preparation including filtering, zero handling, and normalization is on the whole similar to that of association networks. The estimated dissimilarity matrix with entries $d_{kl}$ expressing the dissimilarity between two samples $k$ and $l$ is a $n \times n$ matrix with entries in [0,1] or $[0, \infty)$, depending on the dissimilarity measure. In the latter case, the dissimilarities are scaled to [0,1] by

$$d_{kl} = \frac{d_{kl} - min(d)}{max(d) - min(d)},$$

where $d$ is the vector with dissimilarities of all pairs of subjects (Han et al., 2011).

Two sparsification methods are available in NetCoMi: (i) specifying a threshold so that subjects with a dissimilarity below this value are connected, and (ii) The k-nearest neighbors (KNN) algorithm, where a vertex $v_i$ is connected to the k vertices with minimum dissimilarity to $v_i$ Marchette (2005).

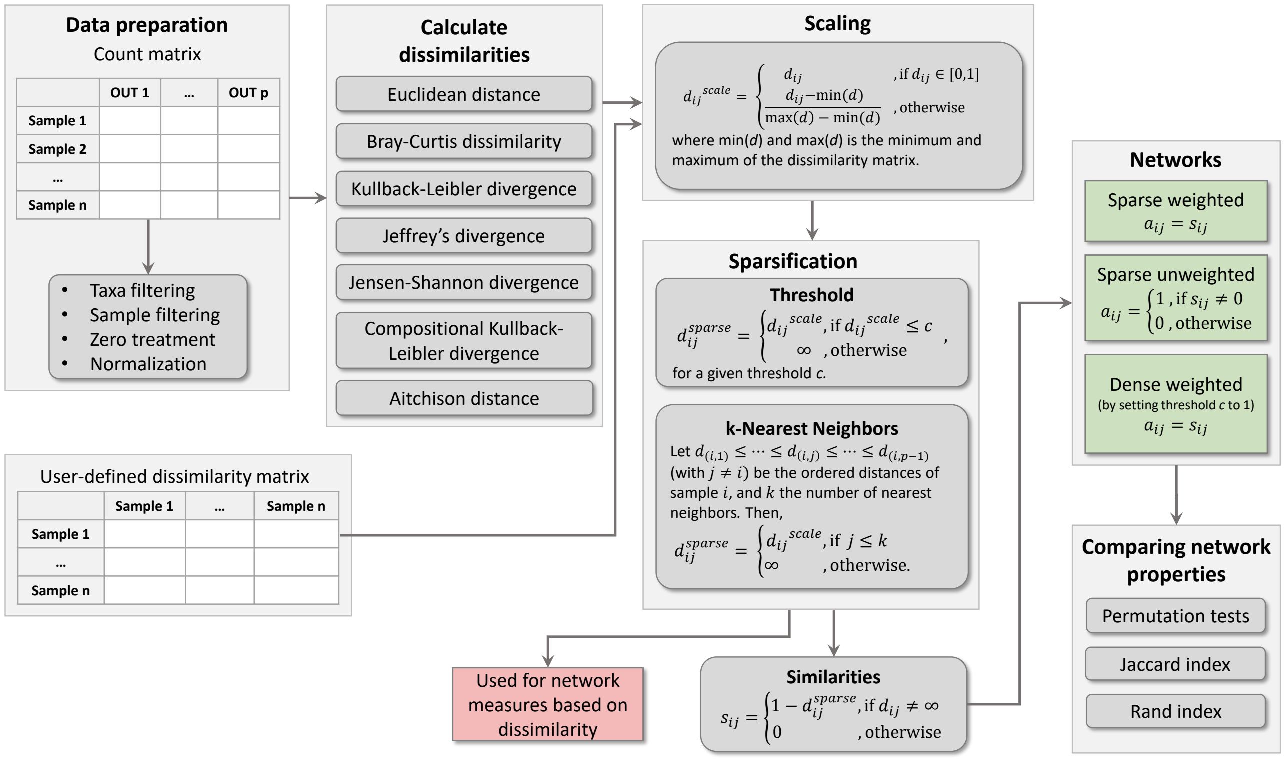

Figure S2: Our proposed workflow for constructing, analyzing, and comparing networks based on dissimilarity between samples. If none of the provided dissimilarity measures should be used, a dissimilarity matrix is accepted as input. If the dissimilarity matrix contains values larger than 1, the values are scaled to [0,1] to facilitate the definition of the threshold (if "threshold" is used as a sparsification method) and to improve the graphical output. The nng() function from the cccd package (Marchette, 2015) is used for constructing a network with k-Nearest Neighbors as a sparsification method. The function implements the approach proposed by Marchette (2005) with an option for constructing a mutual KNN network, where the nodes must be neighbors of each other. The scaled dissimilarities $D(x_k, x_l)^{scale}$ between subjects $k$ and $l$ are used for network measures based on shortest paths. The corresponding similarities (calculated by $1 - D(x_k, x_l)^{scale}$) are used for measures based on connection strength as well as edge weights in the network plot. The network properties of the two networks can finally be compared using the same methods as provided for association networks.

Table S3: Measures of dissimilarity between two compositions $x_k$ and $x_l$ that are implemented in the `NetCoMi` package. These measures are designed for measuring dissimilarities between samples. Thus, in a network based on one of these measures nodes represent subjects or samples instead of taxa.

| Dissimilarity measures | Characteristics |
|---|---|
| **Euclidean distance** <br> $$D_{\text{Eucl}}(x_k, x_l) = \left[\sum_{i=1}^{p}(x_{ki} - x_{li})^2\right]^{\frac{1}{2}}$$ | • symmetric with values in [0, $\infty$) <br> • `R`: `dist()` (`stats` package) |
| **Aitchison distance** (Aitchison, 1992) <br> $$D_{\text{Ait}}(x_k, x_l) = \left[\sum_{i=1}^{p}(\log(\frac{x_{ki}}{g(x_k)}) - \log(\frac{x_{li}}{g(x_l)}))^2\right]^{\frac{1}{2}}$$ <br> ($g(x) = (\prod_{i=1}^{p} x_i)^{\frac{1}{p}}$ is the geometric mean of composition $x$) | • Euclidean distance between $clr$-transformed compositions <br> • symmetric with values in [0, $\infty$) <br> • suitable for application on compositional data |
| **Kullback-Leibler divergence** (KLD) (Kullback and Leibler, 1951) <br> $$D_{\text{KLD}}(P\|\|Q) = \sum_{i=1}^{p} P(x_i) \log\left(\frac{Q(x_i)}{P(x_i)}\right)$$ <br> ($P$ and $Q$ are the probability distribution functions of $x_k$ and $x_l$, respectively) <br> $\Downarrow$ <br> symmetric version (Johnson and Sinanovic, 2001): <br> $$D_{\text{symKLD}}(P, Q) = \frac{1}{2}(D_{\text{KLD}}(P\|\|Q) + D_{\text{KLD}}(Q\|\|P))$$ | • divergence between two probability distribution functions <br> • asymmetric with values in [0, $\infty$) ($D_{\text{KLD}}(P\|\|Q) \neq D_{\text{KLD}}(Q\|\|P)$) <br> • since an asymmetric measure leads to a directed network, only the symmetric version is used in `NetCoMi` <br> • `R`: `KLD()` (`LaplacesDemon` package) |
| **Jeffrey's divergence** (Jeffreys, 1948) <br> $$D_{\text{Jeff}}(P, Q) = D_{\text{KLD}}(P\|\|Q) + D_{\text{KLD}}(Q\|\|P)$$ | • based on KLD <br> • symmetric with values in [0, $\infty$) <br> • `R`: own implementation based on `KLD()` |
| **Jensen-Shannon divergence** (JSD) (Endres and Schindelin, 2003) <br> $$D_{\text{JSD}}(P, Q) = \frac{1}{2}D_{\text{KLD}}(P\|\|M) + \frac{1}{2}D_{\text{KLD}}(Q\|\|M)$$ <br> with $M = \frac{1}{2}(P + Q)$ | • based on KLD <br> • symmetric with values in [0, $\infty$) <br> • `R`: own implementation based on `KLD()` |
| **Compositional Kullback-Leibler divergence (cKLD)** (Martín-Fernández et al., 1999) <br> $$D_{\text{cKLD}}(x_k, x_l) = \frac{p}{2}\log(A(x_k/x_l) \cdot A(x_l/x_k)),$$ <br> where $A(x_k/x_l)$ denotes the arithmetic mean of the vector of ratios $x_k/x_l = \left(\frac{x_{k_1}}{x_{l_1}}, ..., \frac{x_{k_p}}{x_{l_p}}\right)$ | • symmetric with values in [0, $\infty$) <br> • `R`: own implementation <br> • suitable for application on compositional data |
| **Bray-Curtis dissimilarity** (Bray and Curtis, 1957) <br> $$D_{\text{Bray}}(x_k, x_l) = \sum_{i=1}^{p}\frac{\|x_{ki} - x_{li}\|}{x_{ki} + x_{li}}$$ | • symmetric with values in [0,1] <br> • `R`: `vegdist()` (`vegan` package) |

#### 3 Group comparison of network properties

##### 3.1 Permutation tests - Testing centrality measures and global properties for group differences

Centrality measures considered in `NetCoMi` are degree, betweenness, closeness, and eigenvector centrality. To test for each taxon whether these centrality values are significantly different between the two groups, a non-parametric permutation test is performed, which is a common procedure to test graph measures for group differences (Uehara et al., 2014; Hosseini et al., 2012). Own `R` code has been written, which implements the following test procedure.

First, the measures are calculated for the two original adjacency matrices to receive a centrality value for each taxon in both groups. In each repetition, the observed read counts of each subject are then randomly reassigned without replacement to one of the two groups so that the number of subjects within each group equals that of the original data set. For these permuted count data, the association matrices are estimated using the same settings for selecting edges as for the original data. Centrality measures are calculated for the resulting adjacency matrices and finally, the absolute difference between the groups is taken for each measure.

This procedure is repeated many times (1000 by default as suggested by Marozzi (2004)) resulting in a distribution of absolute differences under the null hypothesis that the centrality value is equal in both groups. For each measure, the p-value is calculated as the proportion of permutations in which the absolute difference is greater than or equal to that belonging to the original grouping. A p-value below the desired significance level $\alpha$ means that the observed group difference of the property is significantly higher than expected by chance.

Since $n$ hypothesis tests are conducted simultaneously (where $n$ is the number of nodes), adjusting the permutation p-values for multiple testing is required. As mentioned in Section 2.2 in the main text, two methods for multiple testing adjustment in the case of arbitrary dependencies between the test statistics (which is assumed in our case) are: (1) controlling the local false discovery rate (FDR) as proposed by Efron (2005), and (2) the adaptive Benjamini-Hochberg method (Benjamini and Hochberg, 2000), where the proportion of true null hypotheses is estimated using convex decreasing density estimation according to (Langaas et al., 2005). Both approaches have an increased power compared to the common Benjamini-Yekutieli (Benjamini and Yekutieli, 2001) method, which controls the FDR under dependence as well. Since the number of tests has to be large (several hundred) for method (1), we suggest using method (2) for the permutation tests described in this section.

We implemented the same procedure to test whether the global network properties (average path length, clustering coefficient, modularity, edge and vertex connectivity, and density) are significantly different between the two groups of interest.

##### 3.2 Jaccard's index

In `NetCoMi`, the Jaccard index (Jaccard, 1908) is used for expressing how different the sets of most central nodes are between the two groups. It can be defined via

$$J = \frac{|C_1 \cap C_2|}{|C_1 \cup C_2|} = \frac{C_{12}}{C_1 + C_2 - C_{12}} := \frac{C_{12}}{M},$$

where $C_1$ is the number of species being most central in group 1, $C_2$ the number of nodes that are most central in group 2, and $C_{12}$ the number of species that are most central in both groups (Real and Vargas, 1996).

Following Real and Vargas (1996), we implemented an approach to test whether the observed value for Jaccard's index is significantly different from that expected at random. For this purpose, a distribution of all possible Jaccard values for a specific $M$ can be gained by determining all possible distributions of the $M$ species among the sets $C_1$, $C_2$ or $C_{12}$, and calculating the corresponding Jaccard values (Real, 1999). A table with critical values of the Jaccard index has been published (Real, 1999), which could be used for interpreting the output of `NetCoMi`. However, to facilitate the application of `NetCoMi`, the function for network comparison returns p-values, which are calculated according to the explanations in Real and Vargas (1996). For each Jaccard value $J$, two p-values are returned, which correspond to the hypotheses that $J$ is greater than and smaller than expected at random.

##### 3.3 The adjusted Rand index

The Rand index (Rand, 1971) is calculated as

$$R = \frac{a + b}{\binom{n}{2}},$$

where $n$ is the number of taxa (which is equal in both networks), $a$ is the number of pairs of taxa that are assigned to the same module in both clusterings, and $b$ denotes the number of pairs belonging to different modules in both clusterings.

Qannari et al. (2014) proposed an approach to test whether the adjusted Rand index ($ARI$) is significantly larger than zero, which is the expected value for two random clusterings. They introduce a permutation procedure to generate an empirical distribution of the adjusted Rand index under the null hypothesis that there is no association between the two partitions. For this purpose, a large amount of partition pairs has to be produced with the same structure as the original clustering. More precisely, for each partition the number of groups – and within these groups, the number of elements – should be equal to those of the corresponding original partition. This can be achieved by randomly shuffle the module labels for each partition (Qannari et al., 2014).

Finally, for each generated pair of partitions, the adjusted Rand index is calculated and their arithmetic mean $\overline{ARI}$ and Variance $Var(ARI)$ are calculated. These are used to transform the $ARI$ values into a normalized version:

$$NARI = \frac{ARI - \overline{ARI}}{\sqrt{Var(ARI)}},$$

which is determined for each permutation. Also, the index value for the original partition pair (denoted by $ARI_0$) is transformed into $NARI_0 = \frac{ARI_0 - \overline{ARI}}{\sqrt{Var(ARI)}}$. Based on these results, the p-values are calculated as the proportion of $NARI$ values being larger than the observed value $NARI_0$ (Qannari et al., 2014).

We implemented the procedure in `R` as follows. Clusters are generated via `cuttreeDynamic()` (`dynamicTreeCut` package (Langfelder et al., 2016)). The function returns for each clustering a vector of numerical labels, which assigns each taxon to a cluster. Using `permute()` from the `gtools` package (Warnes et al., 2018) the labels of the two partitions are randomly reassigned and the adjusted Rand index is calculated for the generated clusterings, respectively. Eventually, the normalized index $NARI$ is calculated for all permutations and the original clusterings, and the p-value is calculated as described above. If the determined p-value lies below the desired significance level, the adjusted Rand index for these two clusterings is greater than zero, that is, significantly larger than for two randomly picked partitions.

#### 4 Differential association analysis

From the association measures considered in our work, correlation is the only one for which we have found parametric approaches to test for group differences. Thus, we concentrate on correlations for the moment.

A naive approach to the identify so-called "differential correlations" could be testing if the difference of two estimated correlations is significantly different from zero. This is problematic because the correlation values are only an estimation of the true, unknown correlation coefficient (Fisher, 1915, 1921). Fisher (1915) has shown that shape and variance of their distribution depend on the true correlation value. More precisely, the distribution of the sample correlations $r$ is more skew, the more the true correlation coefficient $\rho$ differs from zero (Fisher, 1921). Thus, the correlations of two populations, which are the two groups of interest in our case, are not directly comparable. To solve this issue, Fisher (1921) suggested transforming the observed values $r$ via $z = \text{arctanh}(r) = \frac{1}{2}\log\frac{1+r}{1-r}$ into Fisher's z-values. These are approximately normally distributed, with standard deviation $\sigma = \frac{1}{\sqrt{n-3}}$ ($n$ denotes the sample size) and thus are suitable for hypothesis testing.

##### 4.1 Fisher's z-test

Fisher's z-test (Fisher, 1992) is a common parametric approach to test the hypothesis $H_0 : \rho_A = \rho_B$ vs. $H_1 : \rho_A \neq \rho_B$ for independent groups $A$ and $B$. First, the sample correlations $r_A$ and $r_B$ are transformed into Fisher's z-values $z_A$ and $z_B$. The aforementioned null hypothesis can be rewritten to $H_0 : \rho_A - \rho_B = 0$, which can be tested using the test statistic

$$Z = \frac{(z_a - z_B)}{\sqrt{\frac{1}{n_A-3} + \frac{1}{n_B-3}}},$$

where $n_A$ and $n_B$ denote the sample size of the two groups. Under the null hypothesis, the Z statistic is standard normal distributed so that standard p-values can be calculated via $p = 2(1 - \mathbb{P}(Z \leq |z|))$.

We use our own `R` implementation for testing whether the correlation coefficients are significantly different between the groups. For multiple testing adjustment, all methods provided by `p.adjust()` (`stats` package) as well as a method for controlling the local false discovery rate provided by the `fdrtool` package (Klaus and Strimmer., 2015) are available in `NetCoMi`.

##### 4.2 Non-parametric tests

As an alternative to parametric tests, a nonparametric resampling-based procedure has been added to `NetCoMi`. The idea for this approach is adapted from Gill et al. (2010, 2014), where taxa in biological networks are tested for differential association between two experimental conditions. Their approaches enable testing whether a single gene is differentially associated with all other genes or whether a set of genes is differentially associated in the two networks. We adopted the procedures to implement a non-parametric approach to test if a pair of taxa is differentially associated, which is a special case of their methods. Own `R` code has been written to make the nonparametric test procedure available in `NetCoMi`.

The authors introduce so-called "connectivity scores" (denoted by $s$ in the following), which represent different types of association (e.g. correlation) between two variables $i$ and $j$. To test whether these variables are differentially associated between networks $A$ and $B$, the distance

$$d(s_{ij}^A, s_{ij}^B) = |s_{ij}^A - s_{ij}^B|$$

is calculated for the observed data first. Then, the two count matrices are combined to a $(m_A + m_B) \times n$ matrix $X$ containing the read counts for both groups. The row numbers are then randomly shuffled to create a permuted matrix $X^*$, which is split again into two groups so that the first $m_A$ rows are assigned to the first group and the last $m_B$ rows to the second. Associations are then determined for the permuted groups and the distances calculated via

$$d^*(s_{ij}^{*A}, s_{ij}^{*B}) = |s_{ij}^{*A} - s_{ij}^{*B}|$$

This procedure is repeated at least $P = 1000$ times, resulting in a vector of $P$ test statistics $d^*$ for each variable. Finally, a p-value for testing the null hypothesis that species $i$ is equally associated in both networks is calculated as the proportion of distances based on the permuted data that are at least as large as the distance belonging to the observed data:

$$p = \frac{\sum \mathbb{1}\{D(s_{ij}^{*A}, s_{ij}^{*B}) \geq D(s_{ij}^A, s_{ij}^B)\}}{P}.$$

Since several hypotheses are tested simultaneously, multiple testing correction is needed for these nonparametric tests as well. The same adjustment methods as for the parametric tests are available in `NetCoMi`.

P-values arising from permutation tests are problematic because they may be exactly zero and a p-value equal to zero remains significant after adjustment for multiple testing. Furthermore, these p-values usually lead to a positively biased type I error rate (Phipson and Smyth, 2010). To overcome this issue, we use the approach proposed by Phipson and Smyth (2010) to estimate exact p-values, which is implemented in R via the `permp()` function (`statmod` package).

Gill et al. (2010) examined the goodness of their approach via a simulation study for a selection of connectivity scores. In that study, correlations, partial correlations as well as connectivity scores based on partial least squares are suggested as appropriate connectivity measures, for which their approaches are supposed to "work well in most circumstances"(Gill et al., 2010). Due to these results, we implemented the test procedure for all association measures (for measuring associations between taxa) available in `NetCoMi`.

The question might arise whether Fisher's z-values could be used as the basis for these tests instead of correlations. This has been done, for instance, in (McKenzie et al., 2016) for conducting differential correlation analysis. In Pesarin (2001) is stated that the power of permutation tests can be improved if based on an – at least approximately – symmetric data distribution. Since Fisher's z-values are approximately normal, permutation tests based on the z-transformed correlations may have a higher power to identify significantly differential correlated taxa. Thus, if a correlation coefficient is used as the association measure in `NetCoMi`, Fisher's z-values can also be used as the basis for the permutation tests instead of the estimated correlations themselves.

##### 4.3 Discordant method

The "Discordant method" (Siska et al., 2016), for which also an `R` package named `discordant` (Siska and Kechris, 2019) exists, is based on mixture models instead of hypothesis testing and is illustrated in Figure S3. Since this approach is based on Fisher's z-transformation, it is only applicable to correlations.

###### 4.3.1 Theoretical Background

Starting from the two correlation matrices belonging to group 1 and group 2, the goal is to assign each pair of the corresponding correlations $r_1$ and $r_2$ to one of the nine classes shown in Figure S3A. This $3 \times 3$ matrix describes all possible correlation scenarios between the groups that are considered by a mixture model. Note that this representation of the matrix is analog to the R code used in the `discordant` package (Siska and Kechris, 2019) (within the illustration of the algorithm pipeline shown in Siska et al. (2016) the groups are interchanged).

The approach is as follows: First, the sample correlations are transformed into Fisher's z-values via $z = \frac{1}{2}\log\frac{1+r}{1-r}$. A mixture model is then created based on these values, which is illustrated in Figure S3B. For each group, the Fisher z-values belong to one of the classes: 0, -, + (denominating the three possibilities: no correlation, negative, or positive correlation between two variables). The z-values within a class are assumed to be normally distributed with mean $\mu$ and variance $\sigma^2$ in group 1 and mean $\eta$ and variance $\tau^2$ in group 2. More precisely, correlations belonging to class "0" are distributed around 0, those in class "-" around an unknown negative mean $\mu_-$ and in class "+" around an unknown positive mean $\mu_+$. For a data set with $n$ variables (or taxa), there are also $n$ variable pairs $k = (z_1, z_2)$ and since the classes (0,+,-) are assigned to single z-values, the paired values belong to one of the classes 1 to 9 of the class matrix in Figure S3A.

If the parameters were known, the class memberships could be easily computed based on the posterior probability that a variable pair belongs to a certain class. On the other hand, if the posteriors were known, a variable pair could be assigned to the class with the highest probability and the parameters would be determined via maximum likelihood estimation.

To solve this issue, an expectation-maximization (EM) algorithm is used, where the two steps are executed iteratively by fixing one component (posterior, or the parameters) and solving for the other.

The density of a pair of z-values is needed for the algorithm, which is defined as follows: Given that $z_1$ belongs to class $i$ and $z_2$ belongs to class $j$ where $i$ and $j$ can take the values 0 (class 0), 1 (class –), or 2 (class +), respectively, the density for the pair of z-values is expressed by:

$$f(z_1, z_2) = \sum_{i=0}^{2} \sum_{j=0}^{2} (\pi_{ij}\ \phi_{\mu_i,\sigma_i^2}(z_1)\ \phi_{\eta_j,\tau_j^2}(z_2))^{\mathbb{1}\{w_{ij}=1\}}.$$

Therein, $\phi_{\mu_i,\sigma_i^2}$ and $\phi_{\eta_j,\tau_j^2}(z_2)$ denote the normal probability distribution functions for group 1 and group 2, respectively, and $\pi_{ij}$ is the proportion of variables with $z_1$ belonging to class $i$ and $z_2$ belonging to class $j$. The membership of the classes 1 to 9 representing all possible correlation scenarios is represented by $w_{ij}$. For example, if a variable pair belongs to class 7, then $w_{2\,0} = 1$ and $w_{ij} = 0$ for $i \neq 2, j \neq 0$. The corresponding likelihood is defined as

$$L(\theta) = \prod_{p=1}^{n} f(z_1^p, z_2^p),$$

with $\theta$ denoting the set of parameters $\{\pi_{ij}, \mu_i, \sigma_i, \eta_j, \tau_j, \}$ for $i, j = 0, 1, 2$ (Siska et al., 2016).

The EM-algorithm is then as follows. First, an initial class membership is needed to determine the means and variances of the classes in both groups. Within the R function, the empirical standard deviations of z-values $s_1$ and $s_2$ are calculated in each group for this purpose. Based on these quantities, all variables with $-s_1 \leq z_1 \leq s_1$ or $s_2 \leq z_2 \leq s_2$, respectively, are assigned to class "0", those with a z-value smaller than the standard deviation to class "-" and those with a larger z-value to class "+". Means and variances are calculated for the resulting classes and used as initial parameters for the algorithm.

In the E-step, for each variable pair $k$, the posterior probabilities are determined for all nine classes (expressed by $i$ and $j$, respectively) via

$$q_{ij}^r(k) = \mathbb{P}(w_{ij}(k) = 1\ |\theta^{r-1}, z_1, z_2),$$

where $r$ is the current iteration number and $\theta^{r-1}$ the parameters of the mixture model resulting from the previous iteration step. $q_{ij}^r$ is the updated posterior probability that a variable pair $k$ belongs to class $(i, j)$ in iteration $r$. Based on these probabilities, the parameters are updated within the M-step via maximum likelihood estimation. The corresponding formulas for re-estimating the parameters can be found in the supplementary material of Siska et al. (2016). These two steps are repeated until convergence of the algorithm leading to posterior probabilities $q_{ij}^r(k)$.

The term of interest, however, is not the probability for one of the nine classes, but the probability that a pair of taxa $k$ is differentially correlated between the classes:

$$p(DC_k) = \sum_{i \neq j} q_{ij}^r(k),$$

which is the sum of posterior probabilities for the off-diagonal classes in the class matrix (Figure S3A). The authors do not suggest a certain threshold above which the taxa are considered as differentially correlated. In `NetCoMi`, we use a default value of 0.8, so that taxa are identified as differentially correlated if the sum of probabilities for the off-diagonal classes is greater than or equal to 0.8.

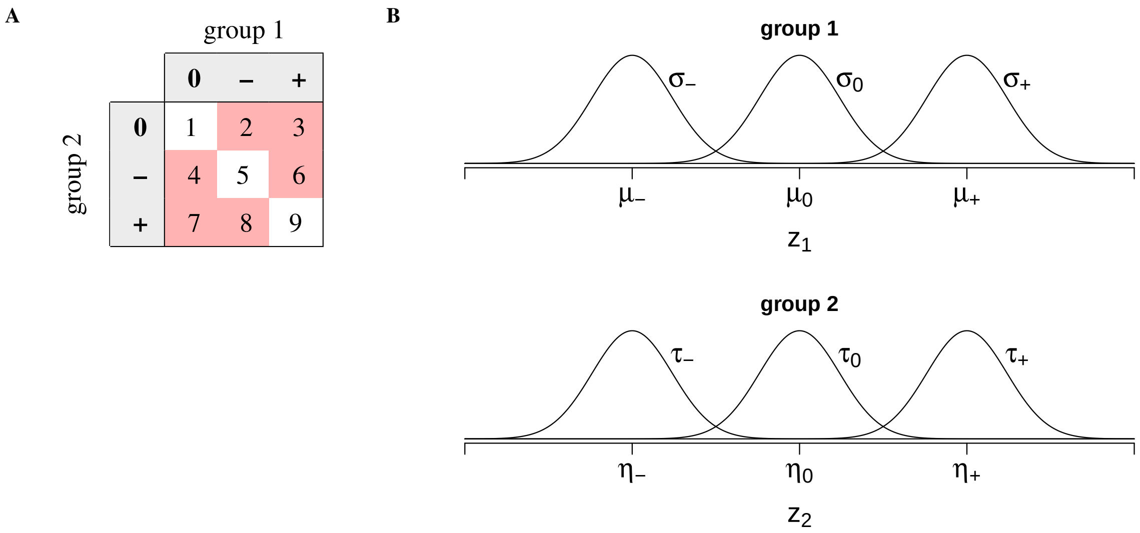

Figure S3: Class matrix and mixture model explaining the Discordant method, which is based on parts of the Discordant algorithm pipeline published in Siska et al. (2016). **A**Class matrix describing all possible correlation scenarios considered in the Discordant approach. Group 1 and group 2 are the two subsets of interest into which the count matrix has been divided. Class 4 means, for example, that two variables are not correlated in group 1 and negatively correlated in group 2. Variable pairs belonging to the off-diagonal classes (colored in red) are considered as differentially correlated. **B** Mixture model with three classes: 0, – and +, for group 1 and group 2 respectively. The z-values within a class are assumed to be normally distributed with mean $\mu$ and variance $\sigma^2$ in group 1 and mean $\eta$ and variance $\tau^2$ in group 2.

##### 4.3.2 Comparison to other methods

Siska et al. (2016) compared their approach to existing methods via a simulation study and additionally by applying the methods to real data for evaluating the performance of the Discordant method. The compared methods for differential correlation analysis were Fisher's z-test, a linear interaction model, and "EBcoexpress" (Dawson et al., 2012) (a bivariate mixture model based on a hierarchical model). P-values were returned by the former and posterior probabilities by EBcoexpress and Discordant, which are not directly comparable. Thus, the variable pairs were ranked by increasing p-values (Fisher's z, EBcoexpress) or decreasing posterior (Discordant). Then, the rankings were compared regarding statistical significance between the methods. Pearson's correlation coefficient was used as correlation measure in each case.

The study concluded that specificity is the same for all methods but Discordant distinguishes from the alternative methods by a higher sensitivity. Thus, more truly differential correlated variable pairs are identified via the Discordant method. Furthermore, the ability of a procedure to predict the paired correlation scenarios expressed by the class matrix were investigated (Siska et al., 2016). It has been shown that all methods have similar performance in identifying the extreme classes where the variables are positively correlated in one group and negative in the other. However, the Discordant method has greater power for detecting differentially correlated variable pairs if the correlation is absent in one of the groups. (Siska et al., 2016).

### 5 Application of NetCoMi to real data from GABRIELA study

#### 5.1 Data preprocessing

Table S4: After denoising the data with QIIME-2 using DADA-2, these preprocessing steps where performed for the mattress dust and nasal swabs data, respectively.

| Preprocessing step | Total number of taxa | Total number of samples |
|---|---|---|
| Mattress dust: | | |
| Original data on OTU level | 61666 | 1044 |
| Remove uncharacterized phyla | 59503 | 1044 |
| Keep only bacteria | 59143 | 1044 |
| Remove Chloroflexi sequences | 56784 | 1044 |
| Remove Chloroplast sequences | 54260 | 1044 |
| Remove mitochondrial sequences | 50533 | 1044 |
| Remove NAs | 45813 | 1044 |
| Assign higher taxonomic rank to uncultured bacteria | 45813 | 1044 |
| Merge down to genus level | 2894 | 1044 |
| Remove samples with less than 1000 reads in total | 2894 | 1022 |
| Remove taxa with less than 1000 reads in total | 707 | 1022 |
| Nasal swabs: | | |
| Original data on OTU level | 46675 | 1127 |
| Remove uncharacterized phyla | 27995 | 1127 |
| Keep only bacteria | 27524 | 1127 |
| Remove Chloroflexi sequences | 26935 | 1127 |
| Remove Chloroplast sequences | 25462 | 1127 |
| Remove mitochondrial sequences | 23937 | 1127 |
| Remove NAs | 20073 | 1127 |
| Assign higher taxonomic rank to uncultured bacteria | 20073 | 1127 |
| Merge down to genus level | 1680 | 1127 |
| Remove samples with less than 1000 reads in total | 1680 | 1033 |
| Remove taxa with less than 1000 reads in total | 467 | 1033 |

##### 5.2 Network properties of the single microbial association network

Table S5: Properties of the network in Figure 4 in the main text. Shown are: a) global network properties, b) frequency table of the clusters, c) the five detected hub nodes, d)-g) centrality values of the ten genera with the highest centrality in decreasing order.

**a)** Global properties

| | |
|---|---|
| average path length | 1.58208 |
| clustering coefficient | 0.28367 |
| modularity | 0.44044 |
| edge density | 0.10586 |
| vertex connectivity | 2 |
| edge connectivity | 2 |

**b)** Clusters

| Name | Frequency |
|---|---|
| 1 | 23 |
| 2 | 31 |
| 3 | 17 |
| 4 | 29 |

**c)** Hub taxa

| |
|---|
| [Eubacterium] coprostanoligenes group |
| Aerococcus |
| Atopostipes |
| Brachybacterium |
| Rikenellaceae RC9 gut group |

**d)** Degree

| **Genus** | **Centrality** |
|---|---|
| [Eubacterium] coprostanoligenes group | 0.22222 |
| Atopostipes | 0.19192 |
| Brachybacterium | 0.18182 |
| Dialister | 0.17172 |
| Sphingomonas | 0.16162 |
| Porphyromonas | 0.16162 |
| Acidiphilium | 0.16162 |
| Aerococcus | 0.16162 |
| Glutamicibacter | 0.16162 |
| Rikenellaceae RC9 gut group | 0.16162 |

**e)** Betweenness centrality

| **Genus** | **Centrality** |
|---|---|
| Porphyromonas | 0.05607 |
| [Eubacterium] coprostanoligenes group | 0.05236 |
| Leuconostoc | 0.04226 |
| Brachybacterium | 0.04061 |
| Sphingomonas | 0.03772 |
| Clostridium sensu stricto 1 | 0.03772 |
| Dialister | 0.03525 |
| Acidiphilium | 0.03504 |
| Blautia | 0.03443 |
| Pseudomonas | 0.03175 |

**f)** Closeness centrality

| **Genus** | **Centrality** |
|---|---|
| [Eubacterium] coprostanoligenes group | 0.87739 |
| Atopostipes | 0.83849 |
| Brachybacterium | 0.83272 |
| Porphyromonas | 0.81466 |
| Rikenellaceae RC9 gut group | 0.81272 |
| Glutamicibacter | 0.80265 |
| Clostridium sensu stricto 1 | 0.80171 |
| Dialister | 0.80013 |
| Kocuria | 0.79544 |
| Prevotella 9 | 0.79531 |

**g)** Eigenvector centrality

| **Genus** | **Centrality** |
|---|---|
| Atopostipes | 1.00000 |
| [Eubacterium] coprostanoligenes group | 0.89872 |
| Brachybacterium | 0.89250 |
| Rikenellaceae RC9 gut group | 0.83003 |
| Aerococcus | 0.79830 |
| Glutamicibacter | 0.77511 |
| Romboutsia | 0.74624 |
| Psychrobacter | 0.70911 |
| Jeotgalicoccus | 0.70841 |
| Prevotella 9 | 0.69081 |

##### 5.3 Microbial association network with "unsigned" distance

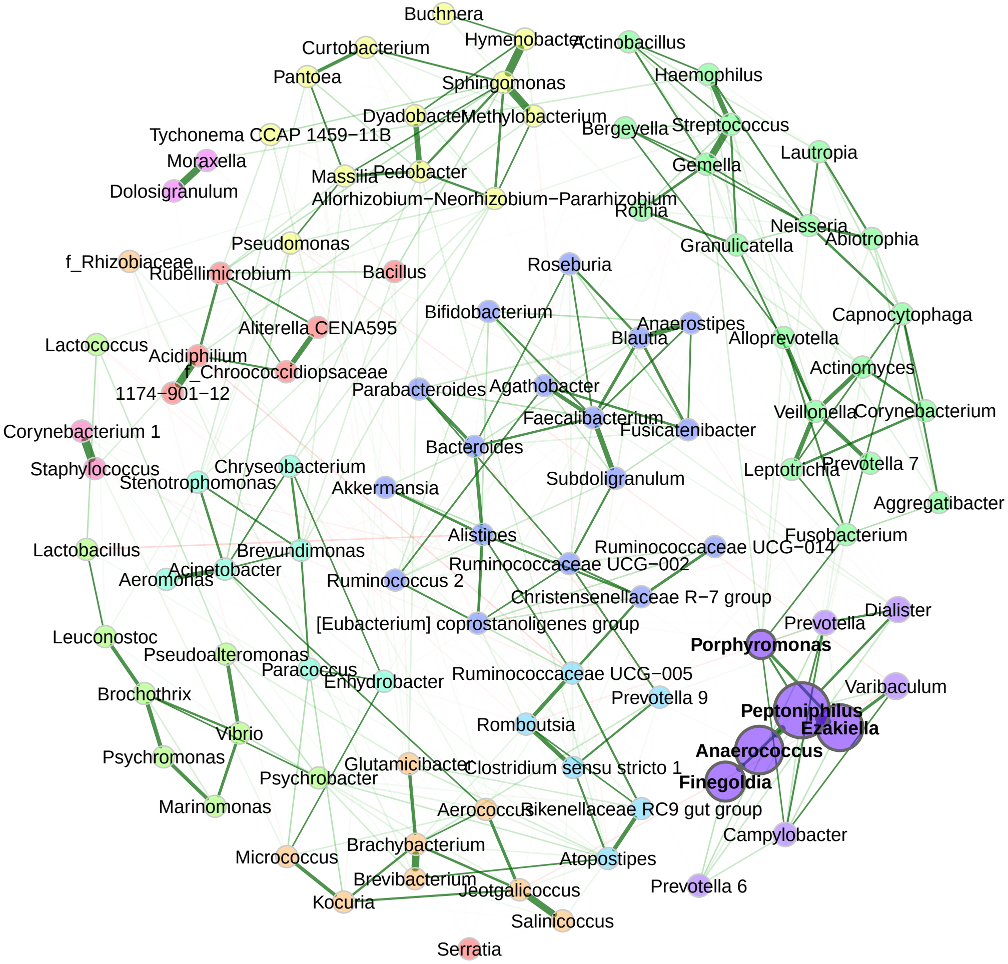

Figure S4: Bacterial associations for the combined data set with samples from Ulm and Munich (with 100 taxa and 1022 samples). The SPRING method Yoon et al. (2019) is used as an association measure. The estimated partial correlations are transformed to dissimilarities via the "unsigned" distance metric and the corresponding similarities are used as edge weights. Eigenvector centrality is used for defining hubs (nodes with a centrality value above the empirical 95% quantile) and scaling node sizes. Hubs are highlighted by bold text and borders. Node colors represent clusters, which are determined using greedy modularity optimization. Green edges correspond to positive correlations and red edges to negative ones.

Table S6: Properties of the network in Figure S4 with "unsigned" distance. Shown are: a) global network properties, b) frequency table of the clusters, c) the five detected hub nodes, d)-g) centrality values of the ten genera with the highest centrality in decreasing order.

**a)** Global properties

| | |
|---|---|
| average path length | 2.34052 |
| clustering coefficient | 0.28367 |
| modularity | 0.38863 |
| edge density | 0.10586 |
| vertex connectivity | 2 |
| edge connectivity | 2 |

**b)** Clusters

| Name | Frequency |
|---|---|
| 1 | 7 |
| 2 | 9 |
| 3 | 12 |
| 4 | 9 |
| 5 | 19 |
| 6 | 7 |
| 7 | 6 |
| 8 | 17 |
| 9 | 10 |
| 10 | 2 |
| 11 | 2 |

**c)** Hubs

| |
|---|
| Anaerococcus |
| Ezakiella |
| Finegoldia |
| Peptoniphilus |
| Porphyromonas |

**d)** Degree

| **Genus** | **Centrality** |
|---|---|
| [Eubacterium] coprostanoligenes group | 0.22222 |
| Atopostipes | 0.19192 |
| Brachybacterium | 0.18182 |
| Dialister | 0.17172 |
| Sphingomonas | 0.16162 |
| Porphyromonas | 0.16162 |
| Acidiphilium | 0.16162 |
| Aerococcus | 0.16162 |
| Glutamicibacter | 0.16162 |
| Rikenellaceae RC9 gut group | 0.16162 |

**e)** Betweenness centrality

| **Genus** | **Centrality** |
|---|---|
| Porphyromonas | 0.05607 |
| [Eubacterium] coprostanoligenes group | 0.05483 |
| Leuconostoc | 0.04082 |
| Brachybacterium | 0.03979 |
| Sphingomonas | 0.03793 |
| Clostridium sensu stricto 1 | 0.03772 |
| Acidiphilium | 0.03711 |
| Blautia | 0.03649 |
| Pseudomonas | 0.03546 |
| Dialister | 0.03381 |

**f)** Closeness centrality

| **Genus** | **Centrality** |
|---|---|
| [Eubacterium] coprostanoligenes group | 0.60208 |
| Atopostipes | 0.57105 |
| Brachybacterium | 0.56016 |
| Rikenellaceae RC9 gut group | 0.55321 |
| Porphyromonas | 0.55316 |
| Glutamicibacter | 0.54612 |
| Dialister | 0.54492 |
| Prevotella 9 | 0.54201 |
| Prevotella | 0.54125 |
| Kocuria | 0.54088 |

**g)** Eigenvector centrality

| **Genus** | **Centrality** |
|---|---|
| Peptoniphilus | 1.00000 |
| Anaerococcus | 0.77458 |
| Ezakiella | 0.72303 |
| Finegoldia | 0.51386 |
| Porphyromonas | 0.19426 |
| Varibaculum | 0.14364 |
| Dialister | 0.11871 |
| Prevotella | 0.08406 |
| Campylobacter | 0.06359 |
| Prevotella 6 | 0.03558 |

##### 5.4 Additional results for network comparison using SPRING as association measure

Table S7: Properties of the networks constructed for the two study centers Munich and Ulm shown in Figure 4 in the main text. Shown are: a) Frequency table of clusters in the Munich network. b) Frequency table of clusters in the Ulm network. c) Detected hub nodes in both groups. d)-g) Centrality values of the genera with the highest centrality in decreasing order. The upper part of the table contains the five genera with the highest centrality in Munich and the lower part those with the highest centrality in Ulm, respectively. Thus, a genus can occur twice in the same table.

**a)** Clusters at Munich

| Name | Frequency |
|---|---|
| 1 | 29 |
| 2 | 6 |
| 3 | 19 |
| 4 | 16 |
| 5 | 30 |

**b)** Clusters at Ulm

| Name | Frequency |
|---|---|
| 1 | 36 |
| 2 | 16 |
| 3 | 17 |
| 4 | 29 |
| 5 | 2 |

**c)** Hubs nodes

| Munich | Ulm |
|---|---|
| [Eubacterium] coprostanoligenes group | [Eubacterium] coprostanoligenes group |
| Aerococcus | Atopostipes |
| Atopostipes | Brachybacterium |
| Brachybacterium | Pedobacter |
| Pseudomonas | Rikenellaceae RC9 gut group |

**d)** Degree

| Genus | Munich | Ulm |
|---|---|---|
| **Highest values in the Munich group:** | | |
| Pseudomonas | 0.20202 | 0.13131 |
| Atopostipes | 0.19192 | 0.18182 |
| Aerococcus | 0.18182 | 0.12121 |
| [Eubacterium] coprostanoligenes group | 0.18182 | 0.19192 |
| Brachybacterium | 0.17172 | 0.16162 |
| **Highest values in the Ulm group:** | | |
| [Eubacterium] coprostanoligenes group | 0.18182 | 0.19192 |
| Rikenellaceae RC9 gut group | 0.10101 | 0.19192 |
| Atopostipes | 0.19192 | 0.18182 |
| Pedobacter | 0.10101 | 0.17172 |
| Neisseria | 0.13131 | 0.16162 |

**e)** Betweenness centrality

| Genus | Munich | Ulm |
|---|---|---|
| **Highest values in the Munich group:** | | |
| Clostridium sensu stricto 1 | 0.04968 | 0.02494 |
| Brachybacterium | 0.04205 | 0.0235 |
| Aerococcus | 0.03628 | 0.01134 |
| Dialister | 0.03546 | 0.01917 |
| Corynebacterium 1 | 0.03484 | 0.01216 |
| **Highest values in the Ulm group:** | | |
| Porphyromonas | 0.02123 | 0.04473 |
| Sphingomonas | 0.0334 | 0.03999 |
| [Eubacterium] coprostanoligenes group | 0.03072 | 0.03669 |
| Gemella | 0.01587 | 0.03608 |
| Neisseria | 0.02597 | 0.03546 |

**f)** Closeness centrality

| Genus | Munich | Ulm |
|---|---|---|
| **Highest values in the Munich group:** | | |
| Pseudomonas | 0.84238 | 0.75576 |
| Aerococcus | 0.83892 | 0.77209 |
| Atopostipes | 0.83707 | 0.82352 |
| Brachybacterium | 0.83634 | 0.79409 |
| [Eubacterium] coprostanoligenes group | 0.83111 | 0.83719 |
| **Highest values in the Ulm group:** | | |
| Rikenellaceae RC9 gut group | 0.73494 | 0.84813 |
| [Eubacterium] coprostanoligenes group | 0.83111 | 0.83719 |
| Atopostipes | 0.83707 | 0.82352 |
| Ruminococcaceae UCG-005 | 0.797 | 0.82 |
| Porphyromonas | 0.72098 | 0.80468 |

**g)** Eigenvector centrality

| Genus | Munich | Ulm |
|---|---|---|
| **Highest values in the Munich group:** | | |
| Atopostipes | 1 | 0.9732 |
| Brachybacterium | 0.90898 | 0.86592 |
| [Eubacterium] coprostanoligenes group | 0.89306 | 0.81747 |
| Aerococcus | 0.87499 | 0.70126 |
| Pseudomonas | 0.85217 | 0.41707 |
| **Highest values in the Ulm group:** | | |
| Rikenellaceae RC9 gut group | 0.59459 | 1 |
| Atopostipes | 1 | 0.9732 |
| Pedobacter | 0.5264 | 0.87283 |
| Brachybacterium | 0.90898 | 0.86592 |
| [Eubacterium] coprostanoligenes group | 0.89306 | 0.81747 |

Table S8: Results from testing centralities of the networks in Figure 5 in the main text for group differences (via permutation tests using 1000 permutations). Shown are respectively the computed measure for Munich and Ulm, the absolute difference, and the p-value for testing the null hypothesis $H_0 : |\text{diff}| = 0$. P-values are adjusted for multiple testing using the adaptive Benjamini-Hochberg method Benjamini and Hochberg (2000), where the proportion of true $H_0$ is determined according to Langaas et al. Langaas et al. (2005). For all four measures, the values are normalized to [0,1] as described in Table 2 in the main text.

| Genus | Munich | Ulm | abs. difference | adj. p-value |
|---|---|---|---|---|
| Degree (weighted): | | | | |
| Corynebacterium 1 | 0.162 | 0.071 | 0.091 | 0.199046 |
| Rikenellaceae RC9 gut group | 0.101 | 0.192 | 0.091 | 0.829358 |
| Lactobacillus | 0.121 | 0.051 | 0.071 | 0.829358 |
| Pedobacter | 0.101 | 0.172 | 0.071 | 0.829358 |
| Pseudomonas | 0.202 | 0.131 | 0.071 | 0.996225 |
| Ruminococcaceae UCG-002 | 0.162 | 0.101 | 0.061 | 0.829358 |
| Aerococcus | 0.182 | 0.121 | 0.061 | 0.996225 |
| Tychonema CCAP 1459-11B | 0.071 | 0.121 | 0.051 | 0.996225 |
| Blautia | 0.152 | 0.101 | 0.051 | 0.996225 |
| Agathobacter | 0.091 | 0.040 | 0.051 | 0.829358 |
| Betweenness centrality: | | | | |
| Aerococcus | 0.036 | 0.011 | 0.025 | 0.978332 |
| Clostridium sensu stricto 1 | 0.050 | 0.025 | 0.025 | 0.978332 |
| Porphyromonas | 0.021 | 0.045 | 0.024 | 0.978332 |
| Corynebacterium 1 | 0.035 | 0.012 | 0.023 | 0.978332 |
| Lactobacillus | 0.026 | 0.003 | 0.022 | 0.978332 |
| Glutamicibacter | 0.025 | 0.004 | 0.021 | 0.978332 |
| Gemella | 0.016 | 0.036 | 0.020 | 0.978332 |
| Brachybacterium | 0.042 | 0.024 | 0.019 | 0.978332 |
| Haemophilus | 0.031 | 0.013 | 0.017 | 0.978332 |
| Dialister | 0.035 | 0.019 | 0.016 | 0.978332 |
| Closeness centrality: | | | | |
| Corynebacterium 1 | 0.825 | 0.690 | 0.135 | 0.884116 |
| Psychromonas | 0.472 | 0.604 | 0.131 | 0.884116 |
| Rikenellaceae RC9 gut group | 0.735 | 0.848 | 0.113 | 0.884116 |
| Finegoldia | 0.694 | 0.591 | 0.103 | 0.884116 |
| Tychonema CCAP 1459-11B | 0.646 | 0.739 | 0.093 | 0.884116 |
| Chroococcidiopsaceae | 0.628 | 0.721 | 0.093 | 0.884116 |
| Alloprevotella | 0.657 | 0.749 | 0.092 | 0.884116 |
| Brochothrix | 0.574 | 0.665 | 0.091 | 0.884116 |
| Paracoccus | 0.646 | 0.734 | 0.088 | 0.884116 |
| Pseudomonas | 0.842 | 0.756 | 0.087 | 0.884116 |
| Eigenvector centrality: | | | | |
| Pseudomonas | 0.852 | 0.417 | 0.435 | 0.574582 |
| Rikenellaceae RC9 gut group | 0.595 | 1.000 | 0.405 | 0.574582 |
| Ruminococcaceae UCG-002 | 0.767 | 0.379 | 0.389 | 0.574582 |
| Corynebacterium 1 | 0.670 | 0.290 | 0.379 | 0.574582 |
| Pedobacter | 0.526 | 0.873 | 0.346 | 0.833965 |
| Blautia | 0.710 | 0.368 | 0.342 | 0.833965 |
| Tychonema CCAP 1459-11B | 0.226 | 0.558 | 0.332 | 0.574582 |
| Ruminococcus 2 | 0.694 | 0.370 | 0.325 | 0.833965 |
| Agathobacter | 0.452 | 0.157 | 0.295 | 0.574582 |
| Rhizobiaceae | 0.324 | 0.602 | 0.278 | 0.833965 |

Significance codes: ***: 0.001, **: 0.01, *: 0.05, .: 0.1

Table S9: Abbreviations of node labels used in Figure 5 in the main text.

| Renamed labels | Original labels |
|---|---|
| Streptoco | Streptococcus |
| Staphyloc | Staphylococcus |
| Acinetoba | Acinetobacter |
| Haemophil | Haemophilus |
| Coryne'1 | Corynebacterium 1 |
| Sphingomo | Sphingomonas |
| Bacteroid | Bacteroides |
| Paracoccu | Paracoccus |
| Pseudomon | Pseudomonas |
| Anaerococ | Anaerococcus |
| Porphyrom | Porphyromonas |
| Faecaliba | Faecalibacterium |
| Prevot'9 | Prevotella 9 |
| Peptoniph | Peptoniphilus |
| Lactobaci | Lactobacillus |
| Prevot | Prevotella |
| Neisseria | Neisseria |
| Ezakiella | Ezakiella |
| Enhydroba | Enhydrobacter |
| Moraxella | Moraxella |
| Gemella | Gemella |
| Rothia | Rothia |
| Actinobac | Actinobacillus |
| Lactococc | Lactococcus |
| Micrococc | Micrococcus |
| Alloprevo | Alloprevotella |
| Veillonel | Veillonella |
| Kocuria | Kocuria |
| Methyloba | Methylobacterium |
| Agathobac | Agathobacter |
| Massilia | Massilia |
| Finegoldi | Finegoldia |
| Pedobacte | Pedobacter |
| Actinomyc | Actinomyces |
| Hymenobac | Hymenobacter |
| Pantoea | Pantoea |
| Subdoligr | Subdoligranulum |
| Bifidobac | Bifidobacterium |
| Psychroba | Psychrobacter |
| Allorhizo | Allorhizobium-Neorhizobium-Pararhizobium-Rhizobium |
| Chryseoba | Chryseobacterium |
| Blautia | Blautia |
| Fusobacte | Fusobacterium |
| Jeotgalic | Jeotgalicoccus |
| Marinomon | Marinomonas |
| Coryne | Corynebacterium |
| Prevot'7 | Prevotella 7 |
| Brochothr | Brochothrix |
| Rumino'us | Ruminococcus 2 |
| Brevundim | Brevundimonas |

| Renamed labels | Original labels |
|---|---|
| Serratia | Serratia |
| Leptotric | Leptotrichia |
| Brachybac | Brachybacterium |
| Brevibact | Brevibacterium |
| Psychromo | Psychromonas |
| Clostridi | Clostridium sensu stricto 1 |
| Rumino'ace | Ruminococcaceae UCG-005 |
| Pseudoalt | Pseudoalteromonas |
| Prevot'6 | Prevotella 6 |
| Granulica | Granulicatella |
| Chrooco(F) | Chroococcidiopsaceae(F) |
| Alistipes | Alistipes |
| Varibacul | Varibaculum |
| Bergeyell | Bergeyella |
| Acidiphil | Acidiphilium |
| Romboutsi | Romboutsia |
| Campyloba | Campylobacter |
| 1174-901- | 1174-901-12 |
| Aerococcu | Aerococcus |
| Dolosigra | Dolosigranulum |
| Rhizobi(F) | Rhizobiaceae(F) |
| Leuconost | Leuconostoc |
| Vibrio | Vibrio |
| Lautropia | Lautropia |
| Dialister | Dialister |
| [Eubacter] | [Eubacterium] coprostanoligenes group |
| Stenotrop | Stenotrophomonas |
| Roseburia | Roseburia |
| Aggregati | Aggregatibacter |
| Bacillus | Bacillus |
| Christens | Christensenellaceae R-7 group |
| Parabacte | Parabacteroides |
| Rumino'ace | Ruminococcaceae UCG-014 |
| Rubellimi | Rubellimicrobium |
| Curtobact | Curtobacterium |
| Atopostip | Atopostipes |
| Glutamici | Glutamicibacter |
| Akkermans | Akkermansia |
| Abiotroph | Abiotrophia |
| Buchnera | Buchnera |
| Rumino'ace | Ruminococcaceae UCG-002 |
| Aliterell | Aliterella CENA595 |
| Rikenella | Rikenellaceae RC9 gut group |
| Anaerosti | Anaerostipes |
| Tychonema | Tychonema CCAP 1459-11B |
| Capnocyto | Capnocytophaga |
| Dyadobact | Dyadobacter |
| Aeromonas | Aeromonas |
| Salinicoc | Salinicoccus |
| Fusicaten | Fusicatenibacter |

##### 5.5 Differential network

A *differential network*, where edges exist between differentially associated taxa, is constructed by passing the object returned from `netConstruct()` to `diffnet()`. We use the permutation procedure (see Table 5 in the main text) to test the associations estimated via the SPRING method (Yoon et al., 2019) for differences. For our data, none of the correlations was significantly different between Ulm and Munich after multiple testing correction (by controlling the local false discovery rate (Efron, 2005)) so that no edges exist in the differential network. Note that this might be due to the fact that permutation p-values have a lower limit, which is related to the number of permutations (in our example 0.001 because of 1000 permutations). Thus, a p-value cannot be arbitrarily small, even if for a pair of taxa the observed difference is much higher than the differences arising from the permuted data.

##### 5.6 Comparison of sample similarity networks

Table S10: Properties of the networks constructed for mattress dust and nasal swabs shown in Figure 6 in the main text. Shown are: a) Frequency table of clusters in the Mattress network. b) Frequency table of clusters in the Nose network. c) Detected hub nodes in both groups. d)-g) Centrality values of the genera with the highest centrality in decreasing order. The upper part of the table contains the five genera with the highest centrality in the mattress network and the lower part those with the highest the centrality in the nose network, respectively. Thus, a genus can occur twice in the same table.

**a)** Clusters in the mattress network

| Name | Frequency |
|---|---|
| 1 | 560 |
| 2 | 221 |
| 3 | 198 |

**b)** Clusters in the nose network

| Name | Frequency |
|---|---|
| 1 | 419 |
| 2 | 245 |
| 3 | 315 |

**c)** Hub nodes (subjects)

| Mattress | Nose |
|---|---|
| 1054 | 1069 |
| 1174 | 1082 |
| 1307 | 1271 |
| 1392 | 1360 |
| 1434 | 1454 |
| 1510 | 1687 |
| 1581 | 1689 |
| 1793 | 1777 |
| 1896 | 1933 |
| 2053 | 2048 |

**d)** Degree

| **Subject** | **Mattress** | **Nose** |
|---|---|---|
| **Highest values in the Mattress group:** | | |
| 1174 | 0.20859 | 0.00307 |
| 1434 | 0.16053 | 0.00307 |
| 1307 | 0.15644 | 0.00307 |
| 1510 | 0.15542 | 0.00511 |
| 1896 | 0.14008 | 0.00307 |
| **Highest values in the Nose group:** | | |
| 1069 | 0.00307 | 0.19632 |
| 2048 | 0.00409 | 0.16462 |
| 1268 | 0.00307 | 0.10327 |
| 1360 | 0.00307 | 0.08998 |
| 1787 | 0.00307 | 0.08998 |

**e)** Betweenness centrality

| **Subject** | **Mattress** | **Nose** |
|---|---|---|
| **Highest values in the Mattress group:** | | |
| 1174 | 0.31951 | 0.00001 |
| 1896 | 0.15342 | 0.00013 |
| 2053 | 0.14126 | 0 |
| 1510 | 0.13824 | 0.00012 |
| 1434 | 0.13787 | 0 |
| **Highest values in the Nose group:** | | |
| 2048 | 0 | 0.45619 |
| 1069 | 0 | 0.32614 |
| 1787 | 0 | 0.17234 |
| 1249 | 0.00417 | 0.11410 |
| 1422 | 0 | 0.10124 |

**f)** Closeness centrality

| **Subject** | **Mattress** | **Nose** |
|---|---|---|
| **Highest values in the Mattress group:** | | |
| 1174 | 4.27319 | 1.05886 |
| 1750 | 4.15324 | 1.46256 |
| 1203 | 3.90505 | 1.42563 |
| 1510 | 3.90289 | 1.20292 |
| 1434 | 3.87384 | 1.38911 |
| **Highest values in the Nose group:** | | |
| 1069 | 2.01941 | 2.28632 |
| 2048 | 2.10909 | 2.28023 |
| 1454 | 2.67159 | 2.08319 |
| 1777 | 2.25869 | 2.02704 |
| 1787 | 1.94227 | 2.02443 |

**g)** Eigenvector centrality

| **Subject** | **Mattress** | **Nose** |
|---|---|---|
| **Highest values in the Mattress group:** | | |
| 1174 | 1 | 0.00151 |
| 1434 | 0.89765 | 0.03591 |
| 1510 | 0.863 | 0.00078 |
| 1307 | 0.72712 | 0.0005 |
| 1896 | 0.70185 | 0.00122 |
| **Highest values in the Nose group:** | | |
| 1069 | 0.048 | 1 |
| 1360 | 0.04889 | 0.59326 |
| 2048 | 0.03478 | 0.40901 |
| 1454 | 0.08403 | 0.38686 |
| 1777 | 0.0695 | 0.30589 |

Table S11, S12, and S13 contain results of the network comparison of the subject-to-subject-networks between mattress dust and nasal swabs shown in Figure 6 in the main text. The determined adjusted Rand index is -0.003 with a p-value of 0.382 meaning that the clusterings are not significantly different from two random clusterings.

Table S11: Jaccard index values corresponding to Figure 6. Index values $j$ express the similarity of the sets of most central nodes and also of the sets of hub taxa between the two networks. "Most central" nodes are those with a centrality value above the empirical 75% quantile. Jaccard's index is 0 if the sets are completely different and 1 for exactly equal sets. $P(J \leq j)$ is the probability that Jaccard's index takes for the present total number of taxa a value less than or equal to the calculated index $j$ ($P(J \geq j)$ is defined analogously). Here, the index is significantly close to zero for all measures, meaning that the sets of most central are considerably different between the groups. The hubs are completely different leading to a Jaccard index of 0.

| | $j$ | P($J \leq j$) | P($J \geq j$) |
|---|---|---|---|
| degree | 0.111 | 0.000000 *** | 1 |
| betweenness centr. | 0.140 | 0.000000 *** | 1 |
| closeness centr. | 0.132 | 0.000000 *** | 1 |
| eigenvec. centr. | 0.153 | 0.000000 *** | 1 |
| hub taxa | 0.000 | 0.000301 *** | 1 |
| Significance codes: ***: 0.001, **: 0.01, *: 0.05, .: 0.1 | | | |

Table S12: Results from testing global network metrics of the networks in Figure 6 for group differences (via a permutation procedure using 1000 permutations). Shown are respectively the computed measure for Mattress and Nose, the absolute difference, and the p-value for testing the null hypothesis $H_0 : |\text{diff}| = 0$.

| | Mattress | Nose | abs. difference | p-value |
|---|---|---|---|---|
| Global network measures: | | | | |
| average path length | 0.516 | 0.892 | 0.375 | 0.000999 *** |
| clustering coefficient | 0.029 | 0.055 | 0.026 | 0.004995 *** |
| modularity | 0.396 | 0.529 | 0.132 | 0.011988 *** |
| edge density | 0.006 | 0.006 | 0.000 | 0.864136 |
| Significance codes: ***: 0.001, **: 0.01, *: 0.05, .: 0.1 | | | | |

Table S13: Results from testing centrality measures of the networks in Figure 6 for group differences (via a permutation procedure using 1000 permutations). Shown are respectively the computed measure for Mattress and Nose, the absolute difference, and the p-value for testing the null hypothesis $H_0 : |\text{diff}| = 0$. The p-values are adjusted for multiple testing using the adaptive Benjamini-Hochberg method (Benjamini and Hochberg, 2000), where the proportion of true $H_0$ is determined according to Langaas et al. (Langaas et al., 2005). These results contain 10 subjects with the highest absolute group difference. All measures are normalized to [0,1].

| Subject | Mattress | Nose | abs. difference | adj. p-value |
|---|---|---|---|---|
| Degree (weighted): | | | | |
| 1174 | 0.209 | 0.003 | 0.206 | 0.017504 * |
| 1069 | 0.003 | 0.196 | 0.193 | 0.017504 * |
| 2048 | 0.004 | 0.165 | 0.161 | 0.017504 * |
| 1434 | 0.161 | 0.003 | 0.157 | 0.017504 * |
| 1307 | 0.156 | 0.003 | 0.153 | 0.039188 * |
| 1510 | 0.155 | 0.005 | 0.150 | 0.656391 |
| 1896 | 0.140 | 0.003 | 0.137 | 0.069617 . |
| 2053 | 0.140 | 0.003 | 0.137 | 0.355379 |
| 1793 | 0.122 | 0.003 | 0.119 | 0.317705 |
| 1268 | 0.003 | 0.103 | 0.100 | 0.017504 * |
| Betweenness centrality: | | | | |
| 2048 | 0.000 | 0.456 | 0.456 | 0.018325 * |
| 1069 | 0.000 | 0.326 | 0.326 | 0.018325 * |
| 1174 | 0.320 | 0.000 | 0.320 | 0.029196 * |
| 1787 | 0.000 | 0.172 | 0.172 | 0.018325 * |
| 1896 | 0.153 | 0.000 | 0.153 | 0.381763 |
| 2053 | 0.141 | 0.000 | 0.141 | 0.880649 |
| 1510 | 0.138 | 0.000 | 0.138 | 0.880649 |
| 1434 | 0.138 | 0.000 | 0.138 | 0.207465 |
| 1581 | 0.131 | 0.000 | 0.131 | 0.018325 * |
| 1307 | 0.126 | 0.000 | 0.126 | 0.189892 |
| Closeness centrality: | | | | |
| 1174 | 4.273 | 1.059 | 3.214 | 0.002017 ** |
| 1392 | 3.843 | 0.967 | 2.876 | 0.001345 ** |
| 1084 | 3.783 | 1.070 | 2.714 | 0.004882 ** |
| 1510 | 3.903 | 1.203 | 2.700 | 0.010759 * |
| 1750 | 4.153 | 1.463 | 2.691 | 0.010289 * |
| 1896 | 3.816 | 1.145 | 2.671 | 0.005505 ** |
| 1793 | 3.811 | 1.219 | 2.592 | 0.006678 ** |
| 1307 | 3.728 | 1.144 | 2.584 | 0.004184 ** |
| 1148 | 3.503 | 0.952 | 2.551 | 0.001345 ** |
| 1434 | 3.874 | 1.389 | 2.485 | 0.004184 ** |
| Eigenvector centrality: | | | | |
| 1174 | 1.000 | 0.002 | 0.998 | 0.048389 * |
| 1069 | 0.048 | 1.000 | 0.952 | 0.048389 * |
| 1510 | 0.863 | 0.001 | 0.862 | 0.338618 |
| 1434 | 0.898 | 0.036 | 0.862 | 0.119550 |
| 1307 | 0.727 | 0.000 | 0.727 | 0.154056 |
| 1896 | 0.702 | 0.001 | 0.701 | 0.174201 |
| 1360 | 0.049 | 0.593 | 0.544 | 0.159400 |
| 1793 | 0.538 | 0.003 | 0.535 | 0.239424 |
| 1054 | 0.482 | 0.008 | 0.474 | 0.082953 . |
| 2053 | 0.520 | 0.076 | 0.444 | 0.287656 |

Significance codes: ***: 0.001, **: 0.01, *: 0.05, .: 0.1

#### 6 Existing simulation studies comparing association and dissimilarity measures

Table S14: Exiting simulation studies, where the performance of the association measures considered in our work is evaluated and compared. Publications containing a simulation study, where one or more of the association measures used in our work is considered, are in rows and the association measures are in columns. The measures designed for sequencing data are ordered by publication date.

| | Traditional measures | | | | Developed for sequencing data | | | | | | |
|---|---|---|---|---|---|---|---|---|---|---|---|
| | Pearson | Spearman | clr-Pearson | Bray-Curtis | CCREPE | SparCC | SPIEC-EASI | CCLasso | gCoda | proportionality | SPRING |
| Year of publication | - | - | - | - | 2012 | 2012 | 2015 | 2015 | 2017 | 2017 | 2019 |
| SparCC paper (Friedman and Alm, 2012) | x | x | x | | | x | | | | | |
| SPIEC-EASI paper (Kurtz et al., 2015) | x | | | | x | x | x | | | | |
| CCLasso paper (Fang et al., 2015) | | | | | | x | | x | | | |
| Review by Weiss et al. (Weiss et al., 2016) | x | x | | x | | x | | | | | |
| gCoda paper (Fang et al., 2017) | | | | | | | x | | x | | |
| propr paper (Quinn et al., 2017) | x | | x | | | | | | x | | |
| Review by Röttjers and Faust (Röttjers and Faust, 2018) | | x | | | | x | x | | x | | |
| SPRING paper (Yoon et al., 2019) | x | | | | | x | x | | | | x |
| Review by Hirano and Takemoto (Hirano and Takemoto, 2019) | x | x | | | | x | x | x | | | |

[Marchette 2005] Marchette, David J.: *Random graphs for statistical pattern recognition*. Bd. 565. John Wiley & Sons, 2005

[Marchette 2015] Marchette, David J.: *cccd: Class Cover Catch Digraphs*, 2015. `https://CRAN.R-project.org/package=cccd`. – R package version 1.5

[Marozzi 2004] Marozzi, Marco: Some remarks about the number of permutations one should consider to perform a permutation test. *Statistica* 64: 193–201 (2004)

[Martín-Fernández et al. 1999] Martín-Fernández, Josep A. ; Bren, Mark J. ; Barceló-Vidal, Carles ; Pawlowsky-Glahn, Vera: A measure of difference for compositional data based on measures of divergence. *Lippard, Næss, and Sinding-Larsen* (1999)

[Mason et al. 2009] Mason, Mike J. ; Fan, Guoping ; Plath, Kathrin ; Zhou, Qing ; Horvath, Steve: Signed weighted gene co-expression network analysis of transcriptional regulation in murine embryonic stem cells. *BMC Genomics* 10, Nr. 1: 327 (2009)

[McKenzie et al. 2016] McKenzie, Andrew T. ; Katsyv, Igor ; Song, Won-Min ; Wang, Minghui ; Zhang, Bin: DGCA: A comprehensive R package for Differential Gene Correlation Analysis. *BMC Systems Biology* 10: 106 (2016)

[Pesarin 2001] Pesarin, Fortunato: *Multivariate Permutation Tests: With Applications to Biostatistics*. John Wiley & Sons, New York, 2001

[Phipson and Smyth 2010] Phipson, Belinda ; Smyth, Gordon K.: Permutation P-values should never be zero: calculating exact P-values when permutations are randomly drawn. *Statistical Applications in Genetics and Molecular Biology* 9, Nr. 1 (2010)

[Qannari et al. 2014] Qannari, El M. ; Courcoux, Philippe ; Faye, Pauline: Significance test of the adjusted Rand index. Application to the free sorting task. *Food Quality and Preference* 32: 93–97 (2014)

[Quinn et al. 2017] Quinn, Thomas P. ; Richardson, Mark F. ; Lovell, David ; Crowley, Tamsyn M.: propr: an R-package for Identifying Proportionally Abundant Features Using Compositional Data Analysis. *Scientific reports* 7, Nr. 1: 16252 (2017)

[Rand 1971] Rand, William M.: Objective criteria for the evaluation of clustering methods. *Journal of the American Statistical Association* 66, Nr. 336: 846–850 (1971)

[Real 1999] Real, Raimundo: Tables of significant values of Jaccard's index of similarity. *Miscellania Zoologica* 22: 29–40 (1999)

[Real and Vargas 1996] Real, Raimundo ; Vargas, Juan M.: The Probabilistic Basis of Jaccard's Index of Similarity. *Systematic Biology* 45: 380–385 (1996)

[Röttjers and Faust 2018] Röttjers, Lisa ; Faust, Karoline: From hairballs to hypotheses–biological insights from microbial networks. *FEMS Microbiology Reviews* 42, Nr. 6: 761–780 (2018)

[Siska et al. 2016] Siska, Charlotte ; Bowler, Russell ; Kechris, Katerina: The discordant method: a novel approach for differential correlation. *Bioinformatics* 32: 690–696 (2016)

[Siska and Kechris 2019] Siska, Charlotte ; Kechris, Katerina: *discordant: The Discordant Method: A Novel Approach for Differential Correlation*, 2019. `https://github.com/siskac/discordant`. – R package version 1.6.1

[Uehara et al. 2014] Uehara, Taira ; Yamasaki, Takao ; Okamoto, Tsuyoshi ; Koike, Takahiko ; Kan, Shigeyuki ; Miyauchi, Satoru ; Kira, Jun-ichi ; Tobimatsu, Shozo: Efficiency of a "Small-World" Brain Network Depends on Consciousness Level: A Resting-State fMRI Study. *Cerebral Cortex* 24: 1529–1539 (2014)

[Warnes et al. 2018] Warnes, Gregory R. ; Bolker, Ben ; Lumley, Thomas: *gtools: Various R Programming Tools*, 2018. `https://CRAN.R-project.org/package=gtools`. – R package version 3.8.1

[Weiss et al. 2016] Weiss, Sophie ; Van Treuren, Will ; Lozupone, Catherine ; Faust, Karoline ; Friedman, Jonathan ; Deng, Ye ; Xia, Li C. ; Xu, Zhenjiang Z. ; Ursell, Luke ; Alm, Eric J. et al.: Correlation detection strategies in microbial data sets vary widely in sensitivity and precision. *The ISME Journal* 10, Nr. 7: 1669–1681 (2016)

[Yoon et al. 2018] Yoon, Grace ; Carroll, Raymond J. ; Gaynanova, Irina: *Sparse semiparametric canonical correlation analysis for data of mixed types*. 2018

[Yoon et al. 2019] Yoon, Grace ; Gaynanova, Irina ; Müller, Christian L.: Microbial networks in SPRING-Semi-parametric rank-based correlation and partial correlation estimation for quantitative microbiome data. *Frontiers in Genetics* 10 (2019)

[Zhang and Horvath 2005] Zhang, Bin ; Horvath, Steve: A General Framework for Weighted Gene Co-Expression Network Analysis. *Statistical Applications in Genetics and Molecular Biology* 4: 17 (2005)
